# Kinetic Control of Nuclear-encoded Mitochondrial mRNA Localization and Local Translation

**DOI:** 10.64898/2026.08.22.746448

**Authors:** Surbhi Sharma, Xuemei Wang, Steven Nguyen, Madeline E. Rasband, Trinh T. Tat, Prabha Chupal, Jen-Yun Chang, Eric L. Van Nostrand, Daniel L. Kiss, Aidan I. Brown, Furqan M. Fazal

## Abstract

Most biological processes are dynamic, yet experimental methods predominantly rely on steady-state measurements to investigate their underlying mechanisms. RNA localization is a fundamental aspect of eukaryotic cell organization and is dynamically regulated by cells. While extensively studied in specialized cell types for a limited number of candidate RNAs, the general principles governing dynamic RNA localization at a transcriptome-wide scale remain largely unexplored. Existing transcriptome-wide studies provide only a static snapshot of RNAs residing in specific cellular locales, in part due to the limited availability of tools for probing cellular spatial organization at biologically relevant scales.

Here, we leverage the high spatial (tens of nanometers) and temporal (minute) resolution of APEX-seq to quantitatively measure the dependence of RNA transport on molecular motors at a transcriptome-wide scale in living cells. We conducted these experiments in the context of the localization of mRNAs to the mitochondria, which are essential for cellular function. Our findings indicate that the majority of nuclear-encoded RNAs encoding mitochondrial proteins localize to the outer mitochondrial membrane (OMM) for local translation. We reveal a crucial role of retrograde dynein-based motor transport in RNA localization, demonstrating that its disruption severely impairs RNA targeting to the OMM.

Time-resolved profiling of RNAs at the OMM revealed that localization is an active process, and even a brief disruption of transport for a few minutes results in a dramatic loss of localization. Moreover, we demonstrate that the translation efficiency (TE) of localized RNAs is a critical determinant of RNA localization in the context of motor-driven transport, as RNAs that delocalize following motor-transport perturbations exhibit lower TE. Using our temporal perturbation data, we also developed a spatiotemporal model that utilizes translation kinetics to capture key features of RNA localization dynamics at the OMM. Together, experiments and modeling suggest that the process of local translation at the OMM is kinetically controlled by the cell, and reveal an unappreciated mechanism by which active transport of RNAs enables cells to modulate their translation within minutes through RNA localization control.

Our study demonstrates how simultaneously capturing the kinetics of hundreds of transcripts with minute resolution can uncover general principles of cellular and organelle organization. Together, these experiments and modeling reveal how active transport and translation jointly maintain the OMM-localized transcriptome. More broadly, they identify RNA localization to cellular membranes as a rapidly tunable mechanism for controlling local translation, even in non-polarized cells.

## Introduction

Eukaryotic cells are highly organized entities that spatially separate the fundamental processes of transcription and translation, with transcription occurring in the nucleus and translation in the cytosol. Messenger RNAs (mRNAs) produced in the nucleus are exported to the cytosol, where they can be translated into proteins. Translational control has therefore emerged as a central mode of post-transcriptional regulation, and much work has focused on how translational machinery, including initiation and elongation factors, is regulated, adapts to cellular stress^1^, and becomes dysregulated in disease^2^. However, recent studies have revealed an additional layer of regulation at the level of the mRNA itself, in which access of transcripts to ribosomes is dynamically modulated. In particular, biomolecular condensates such as stress granules^3,4^ and P-bodies^5,6^ can rapidly sequester RNAs and alter translational dynamics on the timescale of minutes, enabling cells to mount effective responses to acute stimuli. For many stresses, including viral infection, cellular responses are required within seconds to minutes^7^, which is much faster than the typical timescales of transcription and nuclear export, which occur over tens of minutes to hours^8^. One mechanism capable of altering translational dynamics on similarly short timescales is the regulated localization of mRNAs. Here, we investigate the timescale of cytosolic mRNA transport to mitochondria and discover that RNA localization is continuously and actively maintained, such that even brief perturbations lasting only minutes are sufficient to disrupt localization. These findings reveal that RNA localization is highly dynamic and can be rapidly regulated.

RNA subcellular localization and its underlying mechanisms have been actively studied since the 1980s^9–11^. Early work in oocytes demonstrated that RNA localization plays an essential role in spatially restricting protein synthesis, exemplified by transcripts such as *oskar*^12^ and *nanos*^13^. More recently, spatial omics technologies, encompassing both sequencing-^14–19^ and imaging-based methods^20^, have revealed that thousands of RNAs localize to defined subcellular compartments, including nuclear subdomains^21,22^ and cytosolic organelles such as endoplasmic reticulum^14,23–25^, mitochondria^14,23^, stress granules^26,27^, P-bodies^28–31^, and the plasma membrane. Collectively, these findings indicate that mRNA localization is widespread rather than exceptional; however, despite its prevalence, the mechanisms that direct most RNAs to their destinations, and the factors that regulate this process, remain largely unresolved.

In mammalian cells, much of our understanding of the extent and mechanisms of RNA subcellular localization derives from studies in neurons^32^, where thousands of RNAs are differentially enriched in the soma, axons, or dendrites. These and related studies have motivated several hypotheses to explain the prevalence of RNA localization. One widely proposed model invokes spatial control of translation, whereby RNAs are translated at the precise locations where their protein products are needed^33,34^. Another emphasizes efficiency, particularly in highly polarized cells such as neurons, where transporting RNAs to distal compartments and locally synthesizing proteins may be more economical than individually transporting proteins over long distances^35^. Both theoretical modeling and experimental data support these ideas. However, RNA localization is also widespread in relatively small and unpolarized cells, such as human embryonic kidney (HEK) cells^14^, suggesting that these explanations are unlikely to fully account for the phenomenon. Moreover, despite decades of research, the mechanisms by which most RNAs are transported to their destinations, the molecular machinery responsible for this transport, and the extent to which it is dynamically regulated remain largely unknown. Although cis-acting sequence elements, or “zipcodes,” have been identified for a small number of transcripts, such as Map2^36^ and *β*-actin^37^, how the majority of RNAs are targeted is unclear. Furthermore, in some contexts, such as at centrosomes^38,39^, RNA localization appears to be driven primarily by co-translational targeting of nascent polypeptides, with the RNA passively accompanying the protein, raising the possibility that RNA localization can, in certain cases, be a consequence rather than a driver of spatial regulation. Collectively, these observations motivated us to revisit the fundamental reasons why most mRNAs are actively transported within cells.

In this study, we ask what fraction of cellular RNAs are actively transported and how rapidly RNA localization can be altered by acutely disrupting transport through the inhibition of the activity of molecular motors. To examine these questions at transcriptome scale in a tractable system, we focused on RNA localization to the OMM, a membrane whose composition has recently been amenable to interrogation through the development of protein^40,41^- and RNA-based proximity-labeling strategies^14^. Mitochondria are composed of more than 1,000 proteins^42^, nearly all of which (∼99%) are encoded by the nuclear genome, with only 13 proteins encoded by the mitochondrial genome itself. Although mitochondria are enclosed by a double membrane that permits the import of proteins from the cytosol via the TOM–TIM translocase complexes, mRNAs generally do not enter the mitochondria. Although ribosomes have been observed on the mitochondrial outer membrane since the 1970s^43,44^, the question of where most nuclear-encoded mitochondrial mRNAs (hereafter called “mitoRNAs”) are translated remained unresolved for decades. In 2019, we addressed this gap in mammalian cells by developing an RNA proximity-labeling approach, APEX-seq^14^, and demonstrated that hundreds of mitoRNAs localize to and are locally translated at the OMM, similar to what had been observed in yeast^45^. These findings have since been independently supported by multiple RNA- and proteomics-based studies, many of which were performed in HEK cells^46–48^.

Here, we leverage APEX-seq and find that a large fraction of RNAs localizing to the OMM are actively transported, and that even brief disruptions lasting only minutes are sufficient to rewire RNA localization profoundly. This work relies on several key features of APEX-seq. First, proximity labeling enables unbiased, transcriptome-wide mapping of RNA localization at the OMM, allowing for integration with publicly available genomic datasets to infer regulatory principles that would be difficult or impossible to extract from live-cell imaging or RNA FISH analyses of a limited number of transcripts. Second, because RNAs are labeled in living cells, APEX-seq captures localization in an endogenous cellular context, minimizing perturbations and purification-associated artifacts. Third, the rapid labeling kinetics of APEX-seq (∼1 minute) permit acute inhibition of motor-based transport over short timescales (3–30 minutes) that are shorter than typical transcriptional responses. This temporal resolution allows us to isolate the proximal effects of disrupted RNA transport in a regime where global translation patterns, RNA abundance, and cellular morphology remain largely unchanged. Using this approach, we establish a strong dependence of OMM-associated RNA localization on the retrograde motor dynein. Moreover, quantitative modeling reveals that translational kinetics contribute to the dynamic control of mitochondrial RNA localization and local protein synthesis. Together, these findings point to an underappreciated principle of RNA biology: RNA localization functions as a rapidly tunable, temporally controlled switch that can be engaged or disengaged on timescales of minutes to modulate translation.

Given that mitochondrial dysfunction contributes to a wide range of diseases^49^, including aging^50^, neurodegeneration, viral infection, cancer, and cardiovascular disease, understanding how translation at the OMM is rapidly rewired in response to stress has broad biological and therapeutic significance. Beyond their canonical role in energy production, mitochondria integrate innate immune signaling^51^, participate in piRNA biogenesis^52^, and engage in extensive physical and functional interactions with other organelles, most notably the endoplasmic reticulum through mitochondria-ER contact sites^53,54^. Our quantitative modeling further suggests that cells exploit RNA localization as a dynamic means to modulate translation of transcripts on the timescale of minutes. This mode of regulation contrasts with prevailing efficiency-based explanations for RNA localization and instead highlights an underappreciated principle: RNA localization can enable swift, reversible control of protein synthesis in response to changing cellular conditions. In this view, RNA localization to the OMM operates analogously to stress-induced condensates, such as stress granules, but is continuously maintained under basal conditions. Together, our findings extend the conceptual framework of extensive mRNA localization from an efficiency-driven phenomenon to a dynamic, temporally-controlled regulatory strategy.

## Results

### APEX-seq Assay to Investigate Acute OMM mRNA Localization Changes

We have previously used APEX-seq-based RNA proximity biotinylation to nominate over a thousand mRNAs at the OMM in mammalian cells, where many are locally translated^14^. By performing APEX-seq with translational inhibitors, we previously identified two ways by which RNAs reach the OMM. In the first way, which we called ribosome-dependent, mRNAs localize to the OMM in a co-translational state, bound to ribosomes, and aided in localization by the N-terminal targeting peptide (i.e. mitochondrial targeting sequence: MTS). In the second way, which we termed RNA-dependent, mRNAs can get to the mitochondrial surface independent of translation, likely guided by specific cis-elements within the RNA sequence that recruit RNA-binding proteins to transport RNAs to the OMM or facilitate their recruitment to the OMM (Figure 1A). These findings relied on our specific subcellular RNA maps in living cells with high spatial (∼20-100 nm^55,56^) and temporal resolution (∼1 minute), which in turn is determined by the chemistry of the proximity labeling reaction. Building on these findings, we reanalyzed our existing APEX-seq data to estimate the fraction of mitoRNAs that localize to the OMM.

**Figure 1.**
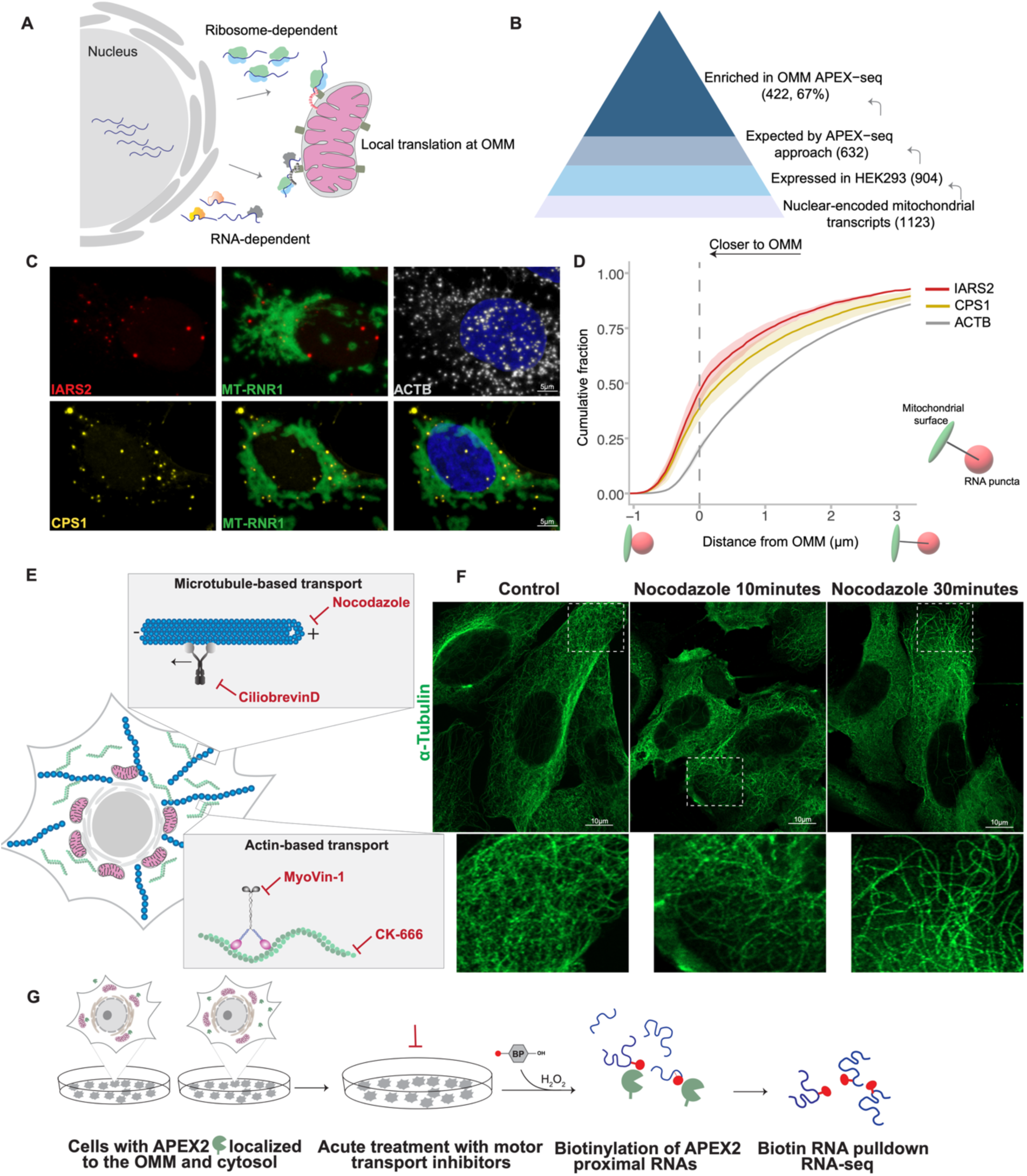
APEX-seq assay to investigate acute RNA localization changes at the OMM in the presence of motor transport inhibitors. (A) Schematic showing the modes of RNA localization to the OMM. (B) A large number of nuclear-derived RNAs coding for mitochondrial proteins localize to the OMM and are detected using APEX-seq approach (∼67%). (C) Representative fluorescence images of maximum-projection views showing mitochondrial network using RNA FISH probes against mitochondrial ribosomal RNA (MT-RNR1), OMM localized candidate RNAs (IARS2 and CPS1), and ACTB in U2OS cells, scale bars-5μm. (D) Cumulative distribution function of the distance between RNA FISH puncta and the mitochondrial surface. Distances were measured in Imaris, and the data shown represent measurements obtained from two biological replicates in U2OS cells. (E) Schematic showing the specific small-molecule inhibitors of the motor-transport used in this study. (F) Immunofluorescence images depicting microtubule (α-tubulin) organization in U2OS cells treated with nocodazole for 10 and 30 minutes, scale bars-10μm. (G) Schematic of the experimental strategy used to profile RNAs at the OMM using APEX-seq in response to motor transport perturbations.

To do so, we first excluded the 13 mitochondrial DNA-encoded genes from the Mitocarta 3.0 database^57^, to estimate the number of mitoRNAs (N = 1123). We then filtered for genes expressed in our cell line by RNA-seq (HEK cells, N = 904) and found 422 of these transcripts localizing to the OMM by APEX-seq. This calculation suggests ∼47% (422/904) of mitoRNAs localize to the OMM. However, this calculation assumes our approach has a 100% sensitivity and can detect every OMM mRNA transcript that is expressed in this cell. If instead we make a reasonable estimate of the detection efficiency of the APEX-seq of ∼70% based on our previous work benchmarking at the well-studied ER membrane (ERM)^14^, which is comparable to other techniques such as biochemical fractionation^24^ that rely on sufficient sequencing coverage to confidently call differentially-expressed genes, we expect to reliably measure the localization of ∼632 (70%) of the 904 mitoRNAs expressed in HEK (Figure 1B). Using this denominator, we thus estimate ∼67% (422/632) of mitoRNAs localize specifically to the OMM by APEX-seq. This estimate is likely an underestimate of the true number, as it excludes transcripts that might also localize to cytosol and other cellular locales that might be missed by our sequencing-based approach. Importantly, this estimate relies on analysis of existing APEX-seq data that captured RNA localization trends at the OMM under treatments with cycloheximide (CHX), puromycin (PUR), and CCCP. Notably, CHX enhances the localization of ribosome-dependent RNAs, while PUR disrupts it. As translational inhibitors dramatically modulate the pool of these transcripts, we proposed that these transcripts localize to the OMM for translation. This estimate of 67%+ is higher than recent estimates of ∼20% from ribosome-profiling studies that study co-translational targeting of ribosomes to OMM^46,47^. We believe this discrepancy can be explained by the fact that our RNA-centric sequencing-based approach is more sensitive and also includes transcripts that localize to the OMM independent of translation (RNA-dependent), but that are indeed translated, including many transcripts for oxidative phosphorylation (OXPHOS) and mitochondrial ribosome components^14^. Figure 1C shows RNA imaging-based (FISH, fluorescent *in situ* hybridization) validation of the localization of representative candidate RNAs (IARS2 and CPS1) at the OMM, measured as the distance (Figure 1D) of these RNA puncta from the mitochondrial surface. Relative to β-actin mRNA (ACTB), these candidate RNAs are enriched at the OMM.

In addition to mitoRNAs, we previously found other types of RNAs in proximity to the OMM, which we separate into two groups: RNAs coding for secretory proteins that we also found at the ERM using ERM-APEX-seq, and other RNAs, including some that are known to be important for mitochondrial function, such as EXTL3^58^ and TSPAN9^59^. The overlap of ERM transcripts at the OMM likely reflects the underlying biology of the ER, which frequently contacts mitochondria through ER-mitochondria junctions^60,61^, and mitochondria are known to divide at ER-mitochondria junctions^62^. In contrast, proximity labeling of the ERM transcriptome using APEX-seq was quite specific for secretory RNAs^14,63^, with no enrichment of mitoRNAs. However, the mechanisms and dynamics of the processes by which all three categories of RNAs (mito, secretory, other) arrive at or near the OMM remain poorly understood. Given that a large number of mitoRNAs localize to the OMM for local translation, and that their mislocalization may impair mitochondrial function, the OMM serves as an ideal model to study if active mechanisms are involved in dictating RNA localization both from a basic mechanism and disease biology perspective.

Cells have a number of motor proteins to processively transport biomolecules, including RNAs, proteins and entire organelles. Most cellular transport occurs along actin and microtubule filaments, aided by molecular motors that recognize, bind, and deliver cargos, including RNAs, to distant locations^64–71^. Previous studies have implicated many motors, such as cytoplasmic dynein^72^, myosin V^73^, kinesin-1^74,75^, and kinesin-3, in the transport of RNAs, both *in vitro* reconstituted systems^76,77^ and in cells. However, most studies so far have focused on the transport of individual RNAs or the cargo preferences of individual motors. We do not know how most RNAs in cells are transported, the kinetics of RNA transport, or whether active transport is dispensable. While these questions are conventionally studied in polarized cells such as neurons^78–80^, intestinal epithelial cells^34,81^ or muscles^82–85^, the role of these processes in small non-polarized cells such as HEK has not been systematically explored. Here, leveraging APEX-seq’s ability to provide unbiased, transcriptome-wide RNA localization data with high spatial and temporal resolution, we systematically dissect how perturbations of active transport disrupt RNA localization to the OMM. In contrast to using traditional imaging-based approaches, comprehensive sequencing enables us to leverage publicly-available datasets to dissect the grammar and features common to transcripts identified, thereby providing broader insights. By pairing our comprehensive sequencing with a spatiotemporal translation-focused model, we find that RNA localization to the OMM relies strongly on active transport, and even a short disruption of transport rewires the local OMM transcriptome to alter translation dynamics.

To study the extent to which individual RNAs rely on active transport, we used specific inhibitors that impair this transport either by disrupting the motor function or the integrity of the cytoskeletal tracks used by the motors (Figure 1E). Disrupting the integrity of the motor tracks impairs bidirectional cargo flow, whereas inhibiting the motor complex impairs directional transport of the targeted motor. For inhibiting actin-based transport, we used CK-666, which prevents the assembly of actin monomers^86^, and MyoVin-1^87^,which inhibits the myosin V motor. Class V myosins are uniquely suited to move cellular cargo over large distances because of their processivity, and have been previously shown to transport RNAs in vitro^88,89^ and in cells^90,91^. For targeting microtubule-based transport we used nocodazole^92^ that disrupts the polymerization dynamics of microtubules (Figure 1F) or ciliobrevin D^93^, a specific inhibitor of the ATPase activity of dynein motor, the sole retrograde transport motor in mammalian cells that moves cargo, including RNAs^65,76,94^, towards the minus end of microtubules.

Our experimental approach (Figure 1G) involves treating cells that have APEX2 targeted to either the OMM or the cytosol with these transport inhibitors. By performing APEX-seq using OMM and cytosol, we can increase the specificity of transcripts recovered at the OMM by performing a ratiometric normalization (OMM/cytosol) using the cytosol transcriptome^14^. We treated cells with inhibitors of motor transport for short durations (between 10 and 30 minutes) to capture early changes in intracellular transport and obtain specific maps of OMM-localized RNAs.

### Inhibition of motor transport disrupts RNA localization to the OMM, as revealed by APEX-seq

Using APEX-seq, we observed an overall decrease in RNA localization at the OMM upon inhibition of motor transport (Figure 2A). Acute treatment with these inhibitors (<30 minutes) combined with the short labeling window of 1 minute employed during APEX2-mediated biotinylation enabled us to investigate dynamic changes in RNA localization without affecting global RNA abundance (Figure 2B), localization of APEX2 tag at the OMM (<u>Figure S1A</u>), mitochondrial positioning and network (<u>Figure S1B</u>), or global translation. Analysis of the APEX-seq libraries confirmed high percentage of mapping, and high correlations between biological replicates (<u>Figure S2A, B</u>). For this study, we investigated more than 1050 mRNAs that we had previously identified at the OMM, with a log_2_foldchange (log_2_FC) (OMM/Cytosol) > 0.75, and FDR < 0.05)^14^.

**Figure 2.**
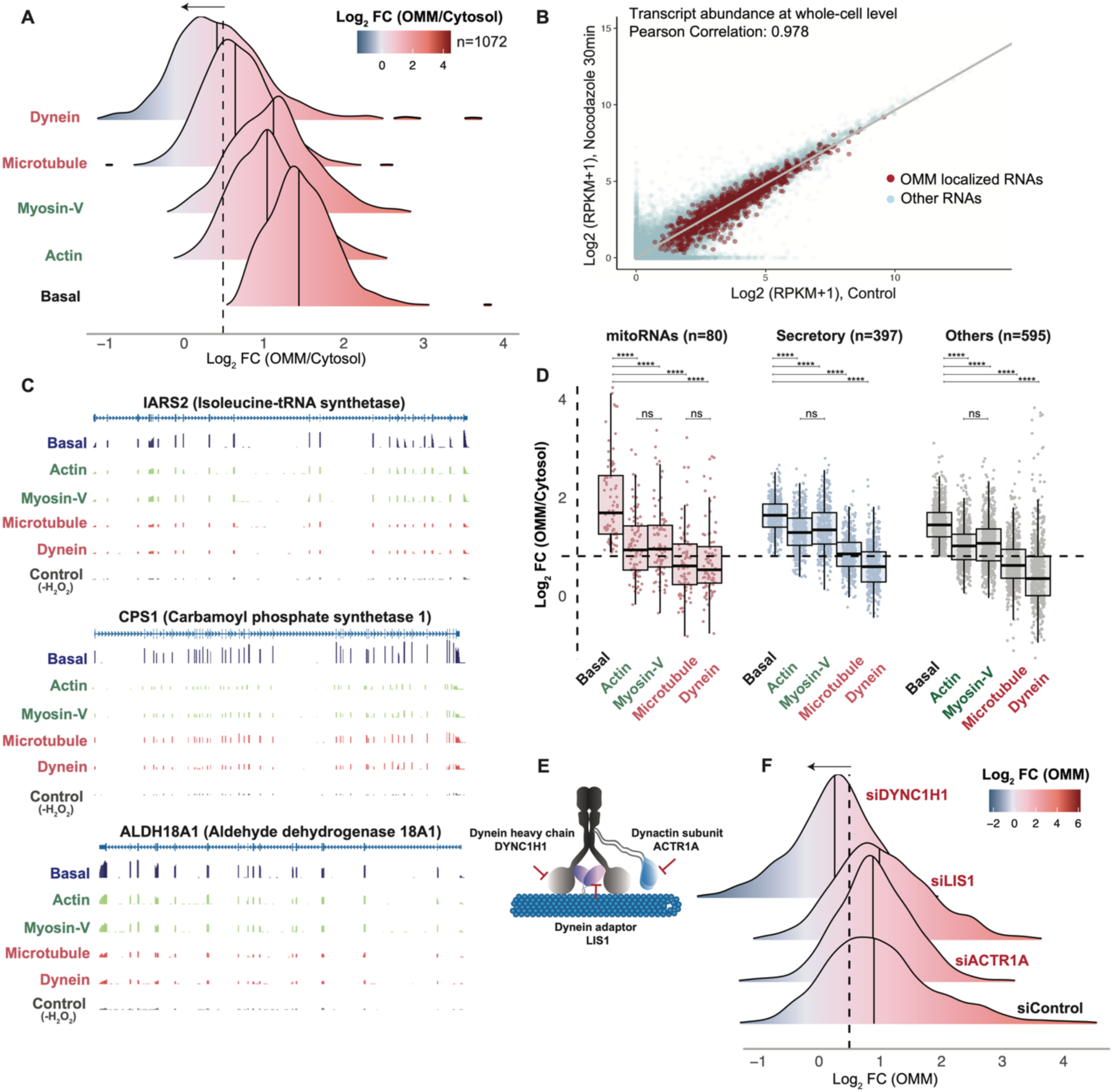
Inhibition of cellular motor transport leads to reduced localization of RNAs at the OMM. (A) Depletion of OMM-localized transcripts upon treatment with inhibitors of actin-based (CK-666: actin; MyoVin-1: Myosin-V) and microtubule-based (nocodazole: microtubule; ciliobrevin D: dynein) transport. The solid vertical lines represent the median log₂ fold change at the OMM, normalized to the cytosol. (B) Correlation plot showing whole-cell transcript abundance in control and nocodazole-treated cells. (C) Representative genome tracks of OMM-localized RNAs (IARS2, CPS1 and ALDH18A1). For each condition, reads were averaged across two replicates; for the control track, reads were averaged across four replicates from both OMM and cytosol conditions. (D) Localization of transcripts in response to motor transport perturbations, with transcripts categorized as mitoRNAs (MitoCarta 3.0), secretory, and others, p-values ****<0.0001 (drug treatment relative to basal) derived using Mann-Whitney U test, ns-not significant. (E) Schematic showing siRNAs used in this study to target components of the dynein motor complex. (F) Depletion of OMM-localized transcripts upon siRNA-mediated knockdown of components of the dynein motor complex. The solid vertical lines represent the median log₂ fold change at the OMM.

Disruption of the actin cytoskeleton or inhibition of the actin-based motor Myosin V resulted in modest reductions in RNA localization, with the log_2_FC of OMM-localized transcripts decreasing from 1.50 ± 0.01 (mean ± SEM) to 1.08 ± 0.01 for actin disruption, and 1.13 ± 0.02 for Myosin V disruption (<u>Figure 2A; S2C</u>). The number of transcripts enriched at OMM (log_2_FC ≥0.75) decreased from 1072 for basal condition to 803 for actin disruption, and 819 for Myosin V disruption. In contrast, inhibition of microtubule-based transport had a dramatic effect on RNA localization, with microtubule network disruption causing the log_2_FC of OMM-localized transcripts to drop to 0.69 ± 0.01 for microtubule disruption. Notably, inhibition of the dynein motor, which drives retrograde transport toward the microtubule minus end, reduced RNA localization the most, with log_2_FC of OMM-localized transcripts falling to 0.48 ± 0.02 (<u>Figure 2A; S2C</u>). Likewise, the number of transcripts remaining after drug treatment was 448 (42%) for microtubule disruption, and only 301 (28%) for dynein disruption. These findings suggest that microtubule-based transport, particularly dynein-mediated retrograde movement, plays a dominant role in directing RNAs to the OMM, with over 70% of transcripts losing their localization upon acute dynein disruption. Figure 2C shows the genome tracks of three representative mitoRNAs, ALDH18A1, CPS1, and IARS2, that display reduced localization upon impairment of cellular motor transport. The data are from polyA-selected RNA, with control tracks generated from the no-labeling control, which we have previously shown is highly correlated (R ∼ 0.95) with bulk RNA-seq data^14^. Thus, mitoRNAs localizing to the OMM are the most sensitive to motor perturbations.

In our previous study^14^, we established three categories of transcripts localizing to the OMM: mitoRNAs (N = 80, under basal conditions), RNAs coding for secretory proteins (typically targeted to and translated at the ERM, N = 397), and other RNAs (N = 595). The overlap of transcriptomes of the ERM and OMM is not surprising, given that the membranes of mitochondria and ER (ERM) have been shown to be tethered by a set of proteins^54^ with the distance between the two membranes ranging from 10-50 nm^53^. Furthermore, cryoET (electron tomography)^61^ and imaging studies^60^ have shown tight coupling between the ER and the OMM^95^, with mitochondria typically dividing at ER-MITO junctions^62^. This close proximity between the two organelles necessitates tight control of the RNA sorting, with several RNAs known to localize to both the ERM and OMM. In the APEX-seq atlas^14^, we observed that approximately 70% of RNAs found at both the ERM and OMM localize to both compartments. Interestingly, disrupting ER RNA localization and local translational machinery has been shown to misdirect such RNAs to the OMM in yeast^96^. Motivated by these observations and to explore potential crosstalk between ERM and OMM RNA localization, we examined how the three categories of OMM-localized RNAs responded differently to drug treatment.

Examining the effect of motor transport on these RNA categories revealed that mitoRNAs, which had the highest average log_2_FC (1.86 ± 0.10) of the OMM-enriched RNAs were also the most affected across all conditions (<u>Figure 2D; S2D</u>), with inhibition of microtubule-based transport showing the strongest impact and a reduction of RNA localization following disruption of the microtubule network and dynein motor (log_2_FC = 0.66 ± 0.08 and 0.63 ± 0.08 respectively). In contrast, inhibition of the microtubule-based transport had a smaller effect on the localization of secretory RNAs, with the corresponding numbers for basal, microtubule-disruption, and dynein disruption being log_2_FC 1.57 ± 0.02, 0.56 ± 0.02, and 0.80 ± 0.02, respectively. Inhibiting actin-based transport, either by disrupting the actin filament or myosin V, disrupted the localization of all categories of transcripts, but to a lesser degree than disrupting microtubule-based transport. Interestingly, mitoRNAs were the most sensitive of the three categories to actin-based perturbation as well, with log_2_FC decreasing from basal (1.86 ± 0.10) for both actin-(1.04 ± 0.08) or myosin V-based (0.97 ±0.08) disruption, respectively.

Intrigued by the finding that acute chemical perturbation of the dynein-based retrograde motor caused the strongest decrease in RNA localization, we further validated our findings by performing siRNA-based genetic knockdown of the dynein motor complex. Dynein is a large multimeric complex with DYNC1H1 (dynein complex heavy chain) as the primary component of the dynein motor^97^. It is assisted by a set of adapter proteins such as LIS1 that help maintain dynein binding to microtubules^98^ and is known to interact with RNA^99^. The dynactin complex proteins (ACTR1A) help tether the cargo to the dynein motor^100^, and are a cofactor for its RNA-transport activity^101,102^. We therefore performed siRNA-based knockdowns of DYNC1H1, LIS1, or ACTR1A and repeated the APEX-seq workflow (<u>Figure 2E; S3A-S3D</u>) to investigate OMM RNA localization. Consistent with our findings from chemical-based perturbation of the dynein motor, knockdown of the dynein motor complex dramatically reduced localization at the OMM (Figure 2F). Relative to an siRNA control experiment, the log_2_FC decreased by 0.84 ± 0.06 upon DYNC1H1 KD (knockdown), supporting the role of dynein in OMM mRNA localization. In contrast, KD of LIS1 and dynactin showed a slight decrease in median localization, but the difference was not significant. Although dynein has been implicated in RNA transport across multiple contexts, ranging from establishing polarity and influencing embryonic development^65,103^, mediating dendritic RNA localization in neurons^104^ to transporting viral particles to the nucleus^105^, our study unveils a novel role for dynein in regulating RNA localization at the OMM.

### Clustering and modeling of time-resolved transport perturbations reveal translational regulation of mRNA localization at OMM

As a sequencing-based RNA localization technique, APEX-seq provides unbiased profiles of mRNAs in different acute treatment conditions. To identify the underlying mechanisms determining why transcripts show differential effects of localization to OMM when different active transport pathways are disrupted, we performed clustering analysis of the OMM-localized RNAs based on the APEX-seq log_2_FC (OMM/cytosol). The analysis finds three distinct groups of transcripts: a group of transcripts somewhat-resistant to motor perturbations (“Stable”, N = 189 genes); a cluster that comprises transcripts lost under all drug perturbations (“Sensitive”, N =379 genes); and the largest cluster comprising transcripts somewhat resistant to actin-but not microtubule-based disruption (“Intermediate”, N =504) (Figure 3A). The transcripts in the stable cluster had the highest basal OMM enrichment (log_2_FC = 1.89 ± 0.05), followed by the intermediate (1.58 ± 0.02) and then the sensitive cluster (1.19 ± 0.02). Notably, upon dynein disruption, the stable cluster which has the highest basal enrichment also showed the least perturbation in enrichment (log_2_FC difference = 0.61 ± 0.07), while both the intermediate and sensitive clusters show similar large localization losses, with the corresponding changes being 1.14 ± 0.03 and 1.05 ± 0.03 respectively. In contrast, upon myosin V transport inhibition, the stable cluster shows no localization loss (log_2_FC difference = 0.02 ± 0.03), intermediate shows a slight loss (difference 0.36 ± 0.01), and the sensitive cluster had the strongest loss (0.57 ± 0.01). Emphasizing this difference in effect upon myosin V disruption, if we classify transcripts with log_2_FC greater than 0.75 as enriched, then not a single transcript in the stable cluster lost localization upon treatment, while 2% were lost in the intermediate cluster (N = 9 of 504), and almost two thirds (64%, N = 243 of 379) were lost in the sensitive cluster. Thus, we can classify our OMM-localized transcripts into three categories that show dramatic differences in localization when treated with the same active transport inhibitors.

**Figure 3.**
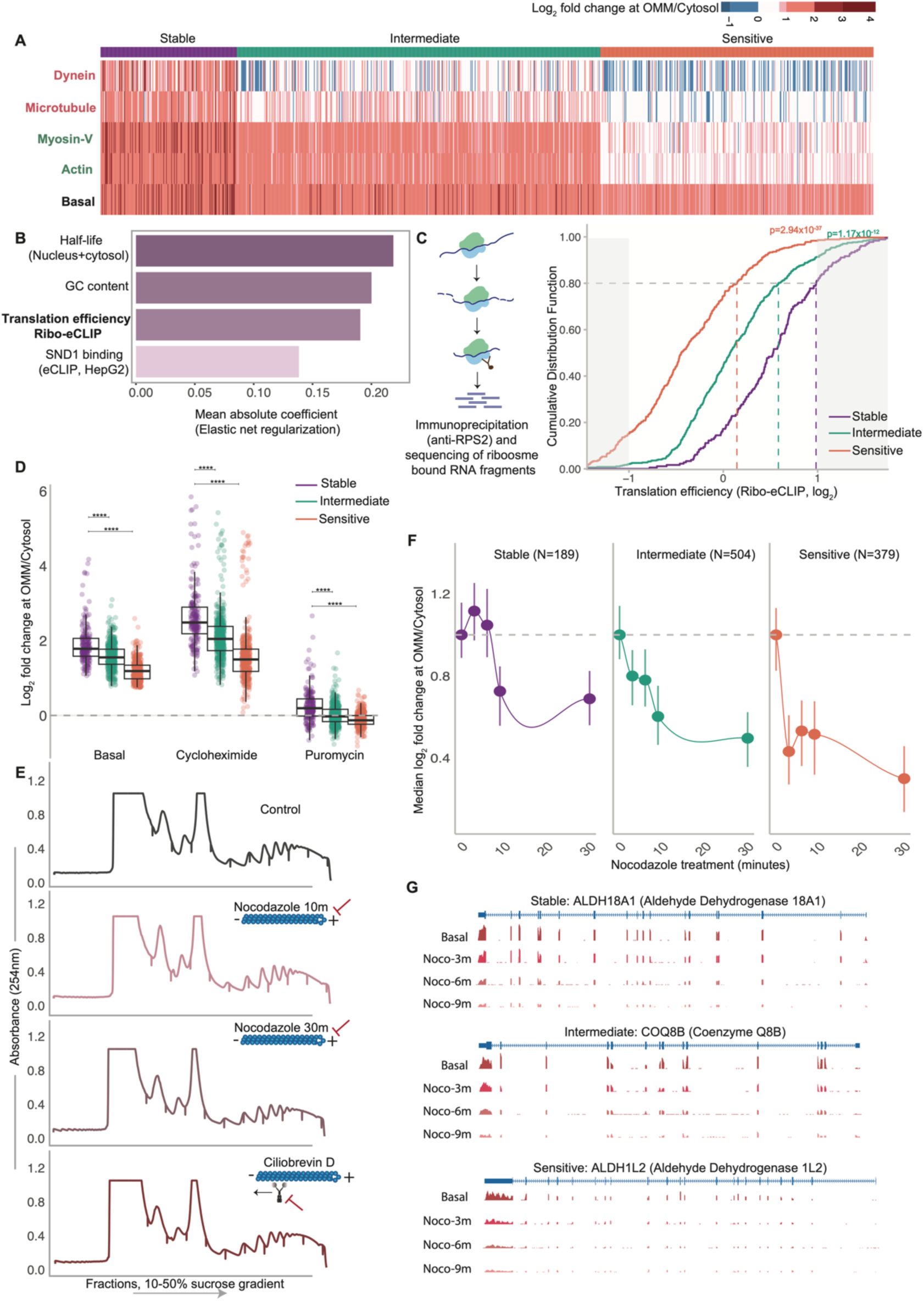
Translation efficiency underlies OMM RNA localization and modulates the response to motor transport perturbations. (A) Heatmap showing log₂ fold changes of OMM-localized transcripts under basal conditions and following treatment with transport inhibitors. Clustering based on fold changes observed under basal and motor-transport perturbations revealed three distinct clusters, categorized by the extent of localization loss at the OMM: Stable (N = 189), Intermediate (N = 504), and Sensitive (N = 379). (B) Top four feature coefficients contributing to cluster identity classification, derived from the elastic net regularization model. (C) Cumulative distribution of the translation efficiency (Ribo-eCLIP) clusterwise, p-values derived from Kolmogorov-Smirnov (KS) tests comparing Stable vs Intermediate and Sensitive clusters. (D) Localization of cluster transcripts at the OMM in the presence of translation inhibitors, cycloheximide and puromycin, p-values ****<0.0001 derived using Mann-Whitney U test, comparing Stable vs Intermediate and Sensitive clusters. (E) Representative polysome profile traces obtained on a 10-50% sucrose gradient from cells treated with inhibitors of microtubule-based transport (nocodazole and ciliobrevin D). (F) Time-course profiling of OMM-localized transcripts across clusters following nocodazole treatment (0, 3, 6, 9, and 30 minutes). Each point represents the cytosol-normalized median OMM localization value at the indicated time point, and bars represent the interquartile range. (G) Representative genome tracks of nuclear-encoded mitochondrial RNAs (MitoCarta 3.0), with one candidate from each cluster. For each condition, reads were averaged across two replicates.

To explain the dramatic changes in outcomes for the three broad categories of transcripts, we investigated a large panel (N = 894) of genetic and molecular features^8,106^ relevant to RNA processing and regulation. These include RNA half-lives, translational efficiency, RNA-protein (enhanced cross-linking and immunoprecipitation, eCLIP^107^) binding datasets, 5′ and 3′ UTR lengths, as well as coding sequence (CDS) and poly(A) tail lengths among others. We evaluated five supervised classification models, including multinomial logistic regression with L2, L1, and elastic net regularization, a random forest classifier, and a histogram-based gradient boosted decision tree model to identify the specific features that are most informative in predicting what transcripts end up in the different clusters. Across all evaluated models, test macro-F1 performance metrics were comparable (0.549-0.582, see <u>Methods</u> section, <u>Figure S4A</u>). However, elastic net regression^108^ that combines lasso (L1^109^) and ridge (L2^110^) regularization to prevent model overfitting, achieved the lowest degree of overfitting among all models (macro-F1_train_ – macro-F1_test_ = 0.067). Based on this, we selected the elastic net model for downstream feature selection, although all models consistently identified the same top three features (half-life, translation efficiency and GC content).

To ensure the consistency of top features identified by the elastic-net model, we performed the model simulation over 300 times and consistently found the same five features (<u>Figure 3B and S4B</u>) that could best explain the observed categories. Among these top five features, four were positively associated with the stable cluster (mean elastic net coefficient >0.1): translational efficiency, half-life, GC content, SND1 binding. As translational efficiency was the most predictive feature and translational control is a key aspect of gene regulation, we decided to focus on its role in RNA localization to mitochondria in subsequent sections.

RNA half-life also emerged as a key predictor. Further investigation of the data revealed that it is not half-life of the transcript in the cytosol, but rather the lifetime of the RNA on the chromatin (<u>Figure S4C</u>^8^) that correlates with the three clusters. These findings are particularly intriguing, as they suggest that nuclear RNA processing can influence localization^111^. This idea is further supported by a recent study^47^ demonstrating that splicing can regulate local translation of mRNAs at the OMM, although the precise mechanisms and the RNA-binding proteins^112^ involved remain poorly understood. Similarly, GC content is another predictive feature, with transcripts in the stable cluster having the highest GC content of 55.2% ± 0.5% (N = 187) and those in the sensitive cluster showing the lowest 46.0% ± 0.4% (N = 378), though again the reasons remain unclear. Among the remaining top predictive features, two were CLIP-related (SND1 and MOV10 binding). SND1 is a core component of the RNA-induced silencing complex (RISC)^113,114^ and plays a critical role in microRNA-mediated regulation, whereas MOV10 functions as an RNA helicase involved in RNA decay^115^. Together, these features highlight multiple layers of post-transcriptional regulation associated with the clusters, although their precise mechanistic contributions will require further investigation.

### Translation and sensitivity to motor perturbations

Translational efficiency is one of the strongest predictive features distinguishing RNAs between the clusters (Figure 3B), with an elastic net coefficient of 0.19. We obtained translation efficiency measurements used in the model by performing a CLIP-based experiment called ribo-eCLIP in HEK293T cells, which quantifies translational efficiency by profiling ribosome-associated RNA fragments relative to total mRNA (Figure 3C) (Chang JY & Van Nostrand EL, *in preparation*). Analysis of the ribo-eCLIP data revealed that RNAs in the stable cluster had the highest TE of 1.53 ± 0.04 (N = 180), followed by 1.19 ± 0.02 (N = 474) for the intermediate cluster and 0.84 ± 0.02 (N =360) for the sensitive cluster. These changes between clusters are quite large and biologically meaningful, especially compared to the whole translatome^116,117,118^. Sensitive-cluster transcripts were significantly enriched for low TE transcripts, with 25.8% (N =93 of 360) having log₂(TE) < −0.75 compared with 14.6% of the translatome (odds ratio = 2.08, Fisher’s exact test, P = 1.5×10⁻⁸). Similarly, stable-cluster transcripts were significantly enriched for high TE transcripts, with 30.0% having log₂(TE) > 0.75 compared with 18.3% of the whole translatome (odds ratio = 1.94, Fisher’s exact test, P = 1.3 × 10⁻⁴). We thus propose that these classes of transcripts respond differently to acute transport inhibition. Specifically, we hypothesize that it is translation-mediated anchoring of transcripts to the OMM that explains why the stable clusters of transcripts remain more resistant to drug treatment.

To validate the role of translational-anchoring in explaining the differences between how transcripts in different clusters respond to perturbations, we analyzed our previous APEX-seq dataset^14^, which profiles the OMM-localized RNA repertoire under translational inhibition using the drugs cycloheximide (CHX) or puromycin (PUR). Specifically, we previously demonstrated that there are two pathways by which transcripts show up at the mitochondria (Figure 1A), either as part of a translating ribosome where the localization is driven by the nascent polypeptide (MTS for mitoRNAs), or through a ribosome/translation-independent mechanism where the RNA localizes to the OMM based on sequence-elements within. The former group of transcripts localization is enhanced upon CHX treatment and disrupted upon PUR treatment, while the latter remains resistant to PUR treatment; these findings have recently been confirmed by a proximity-specific ribosome profiling study^46^. Analysis of these datasets revealed that differences in OMM localization among the clusters became more pronounced in the presence of CHX, which enriches for active-translating mRNAs (Figure 3D). Specifically, the OMM-enrichment log_2_FC differences between the stable and sensitive cluster are highest upon CHX treatment (difference log_2_FC = 1.03 ± 0.07), and least upon PUR treatment (log_2_FC = 0.37 ± 0.04). Additionally, the stable cluster, which already has the highest basal enrichment to OMM, increases localization the most under CHX treatment (log_2_FC increase = 0.76 ± 0.07), and decreases the most under PUR treatment (log_2_FC change = –1.64 ± 0.05). The corresponding numbers for the sensitive cluster, in contrast, are 0.42 ± 0.05 and –1.31 ± 0.03 respectively. Thus, re-analysis of our prior OMM APEX-seq datasets under translational inhibitors confirms translation to be a key factor segmenting these transcripts.

Taken together, our data so far support ribosome-mediated anchoring to the mitochondrial surface as a key feature distinguishing how transcripts respond differently to active-transport inhibition. To rule out other explanations for the data, we explored confounding factors that might explain these findings: changes in RNA stability or RNA decay, global translation changes, OMM-APEX2 tag displacement, and rapid mitochondrial reorganization. For the first alternative explanation, we show in Figure 2B that globally we see no significant differences in RNA levels pre- and post-drug treatment, including the 1000+ OMM-localized transcripts considered in this study. For the second explanation, we performed a polysome profiling experiment using a sucrose gradient to study loading of ribosomes to mRNAs pre- and post-drug treatment. We performed these experiments on cells treated with either microtubule-inhibition (nocodazole) or dynein inhibition (ciliobrevin D), conditions under which we observed the largest decreases in localization. In both conditions, we did not see substantial differences in the polysome fraction at these acute treatments compared to controls (<u>Figure 3E & S4E</u>). As cellular stresses typically induce a conserved response where the translational factor eIF2*α* undergoes phosphorylation^119–122^, followed by the shutdown of translation of many transcripts and sequestration of mRNAs in stress granules, we also checked phosphorylated eIF2α levels by western blot in these conditions. Again, we did not observe induction of phosphorylation in response to the acute treatments used in this study (<u>Figure S4D</u>), thereby supporting the observation that the differences in localization to the OMM are driven by specific differences between the TE of these RNAs, and not due to global translation changes.

To confirm that the OMM-APEX2 tag remains on the OMM upon drug treatment, we performed an immunofluorescence (IF) experiment with and without nocodazole treatment. At the acute times (<30 minutes) we interrogated, we observed no change in localization of APEX2 tag to the OMM (<u>Figure S1A</u>). Lastly, using IF we examined the positioning and morphology of mitochondria upon nocodazole 10-minute or 30-minute treatment. Indeed at 10 minutes nocodazole treatment we observed no appreciable change in mitochondrial morphology or their perinuclear positioning within cells (<u>Figure S1B</u>). At 30 minutes, mitochondria continue to be positioned at the nuclear periphery and their morphology is somewhat preserved. As we have previously shown beyond 30 minutes, such as a 2-hour drug treatment, mitochondria change their morphology, as well as RNA levels (RNA-seq revealed differentially-expressed transcripts)^14^. Taken together, these data support our initial inference that the acute changes in RNA localization to mitochondria upon disruption of active transport are largely driven by two factors: motor, particularly dynein-driven, transport; and ribosome-anchoring at or near the mitochondrial surface.

The acute treatments, lasting 10 to 30 minutes depending on the condition, yielded dramatic changes in RNA localization to the OMM in a timescale when RNA levels, organelle positioning, and global translation remain unaltered. These observations prompted us to investigate whether the kinetics of RNA localization loss might differ between the different clusters. To do so, we reanalyzed our previous APEX-seq time course nocodazole experiments^14^ and studied how transcripts belonging to the different clusters are lost. Time-resolved analysis revealed distinct localization kinetics among the stable, intermediate, and sensitive clusters. Specifically, within a 3-minute perturbation coupled with APEX-seq, we already find the sensitive cluster is lost at the OMM. In contrast, up to about the first 6 minutes, the stable cluster, shows almost no loss of localization as a whole at the OMM (Figure 3F). The intermediate cluster, transcripts are intermediate in their loss, showing a more gradual loss with increasing treatment time (half-life of loss ∼5 minutes). These experiments leveraging APEX-seq to perform unbiased, rapid, transcriptome-wide, measurements of RNA localization in living cells reveal changes in kinetics of transcripts that occur at short (∼1-3-minute) times. In general, when we segment our OMM-localized transcripts into ones that are mitoRNAs vs transcripts co-localizing to ER and MITO (<u>Figure S4F</u>), we find the former are in general more quickly lost than the latter in all clusters. However, across all clusters, stable-cluster transcripts were lost much more slowly than intermediate or sensitive clusters. Although in this study we focus on the aggregate behavior of hundreds of transcripts, review of individual transcripts illustrates these dynamic differences (Figure 3G). For example, the mitoRNA ALDH18A1 remained relatively more stably associated with the OMM for the first few minutes, relative to ALDH1L2 that is lost relatively quickly. The intermediate transcript COQ8B shows progressive loss upon increasing nocodazole treatment time.

Given that the typical translation time (*τ*) of a mRNA is ∼1-3 minutes, we think our data fit well with a ribosome-mediated anchoring model. Specifically, almost all OMM-localizing transcripts require active transport to get them to the mitochondria. However, when this process is disrupted, transcripts with one or a few ribosomes loaded diffuse away after about one round of translation (t ∼ *τ*). In contrast, stable transcripts have multiple ribosomes bound (i.e. high TE) and, as such, do not diffuse away from the mitochondria immediately upon active transport disruption (t > *τ*). However, once all ribosomes have terminated translation, the transcript is no longer in proximity to the mitochondria, presumably because upon perturbation of active transport, diffusion of the transcript away from the mitochondria dominates. Supporting this model, it is notable that the negative-end directed motor dynein, which is known to bring transcripts towards the nucleus^65,76,94,123^, is most important for localization. The mitochondria, being perinuclear in our system, need active transport to keep transcripts near the nucleus for translation. Cells cannot rely on diffusion to get mRNAs to the mitochondria, as a diffusive outcome is a poor outcome in this system since it will distribute the transcript across the volume of the cell, most of which is devoid of mitochondria (<u>Figure S1B</u>).

In summary, our findings suggest active transport is essential for appropriate localization of transcripts to mitochondria, which in our cell type with perinuclear mitochondria, is dominated by dynein. Presumably in other cell types, or in mitochondria in other locations, the kinetics of RNA transport and subsequent translation will likely differ, which in turn may alter the composition of the mitochondria and contribute to mitochondrial heterogeneity within the same cell^124–127^.

### A spatiotemporal model links motor transport and translation to kinetically controlled OMM mRNA local translation

Combining APEX-seq with acute transport perturbations allowed us to quantify transcript loss from the OMM. The data are qualitatively explained by a model in which active transport delivers mRNAs and ribosome-mediated anchoring retains them at mitochondria. However, the fact that transcripts rapidly (within minutes) diffuse away from the mitochondria, and that constant active transport is required to maintain proper mRNA localization to OMM, at first seem to be in contradiction with the general view of mRNA localization mechanisms evolving for efficiency, as has been demonstrated well in neurons^35,128,129^. mRNA localization in neurons provides efficiency gains, as it is less energetically expensive to transport an RNA to its destination and have it translate multiple proteins, rather than moving the individual proteins after translation. But it is also the case that RNA localization is a wide-spread phenomenon, with our previous HEK APEX-seq datasets generating 3000+ localized RNAs from 9 subcellular locations, and numerous studies in neurons^130–134^ or drosophila oocytes^135,136^ finding thousands of localized transcripts. As such, the reasons for RNA localization being so widespread continue to remain an active area of investigation. The requirement to constantly transport mRNAs in our system for correct OMM localization, an active process using energy, motivated us to consider whether the cell might be using RNA localization as a method of translational control. In other words, might localization to the mitochondria be a tunable knob that cells can use to dynamically increase or decrease translation? Our quantitative profiling of RNA loss for hundreds of RNAs on minute (∼minutes) timescale gives us a unique insight into events occurring on a translation-relevant (*τ*) timescale, a regime rarely explored in biological studies at a systems-wide scale, even when those studies that do have kinetic information.

To test these ideas, we developed a quantitative kinetic model (Figure 4A&B) of translation-mediated association with mitochondria, similar to recent work^137^, to better understand how intracellular transport perturbations affect mRNA association with mitochondria. This model describes mediation of mRNA association with the mitochondria through the nascent peptide sequences (MTS for mitochondria) that facilitate import of the mature protein into the mitochondria through the TIM-TOM^138,139^ complex. Thus, mRNAs that have at least one currently-translating ribosome with a binding-competent nascent peptide are anchored to the mitochondria. This model implicitly builds in the fact that mitochondria are perinuclear, so perinuclear RNAs with a binding-competent nascent peptide associate with mitochondria (k_on_) and mitochondrially-associated RNAs that lose all binding-competent nascent peptide dissociate from the mitochondria (k_off_). RNAs leave the perinuclear region for the periphery at rate k_leave_ and return at rate k_return_. Further details of the model are in the <u>Methods</u> and illustrated in <u>Figures 4A and 4B</u>.

**Figure 4.**
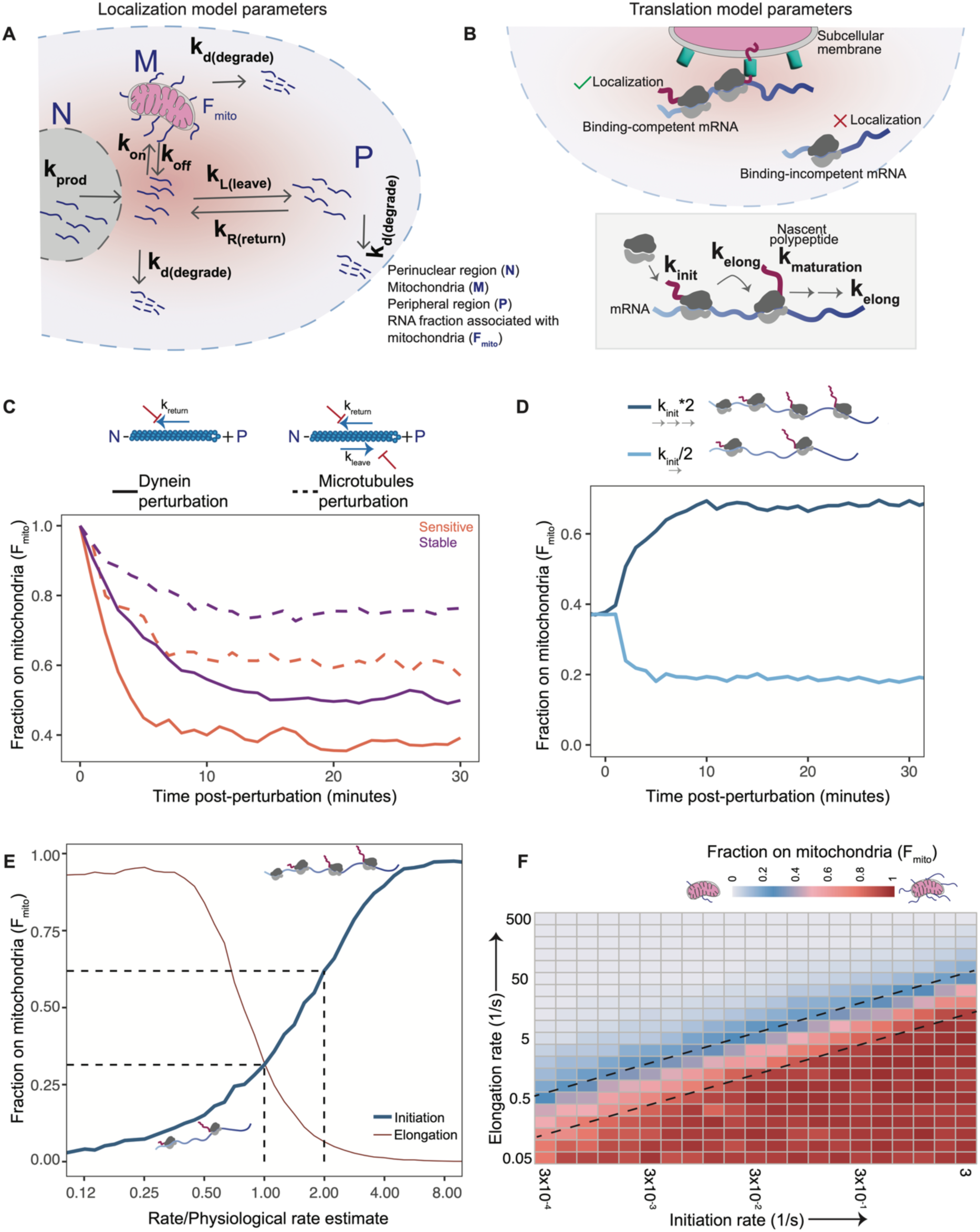
A spatiotemporal model linking motor transport and the translation process in RNA localization. (A) & (B) Schematic of the model coupling translation and cellular localization. The rate constants (k) used for stochastic simulations are shown. (C) Fraction F_mito_ of mRNA associated with mitochondria vs time since transport parameter perturbation. Orange and purple dotted curves show change in F_mito_ (for sensitive and stable clusters respectively) following decrease in k_leave_ by a factor of 2 and k_return_ by a factor of 4, representing perturbation of microtubules by nocodazole. Orange and purple solid curves show changes in F_mito_ (for sensitive and stable clusters respectively) following decrease in k_return_ by a factor of 4, representing perturbation of dynein-driven transport. (D) Fraction F_mito_ of mRNA associated with mitochondria vs time since translation parameter perturbation. Curves show the change in F_mito_ (for an intermediate initiation rate) following a two-fold increase or decrease in the initiation rate, representing cellular changes that alter initiation. (E) Steady-state fraction F_mito_ of mRNA associated with mitochondria vs translation initiation (thick blue curve) and elongation rate (thin red curve). Vertical dashed black lines show where the localization has doubled for a 2-fold decrease in initiation timescale. (F) Steady-state F_mito_ (indicated by color) as translation initiation (horizontal axis) and elongation (vertical axis) rates are varied, representing that much of the possible range of mitochondrial localization is covered by a narrow band of elongation and initiation timescales (dashed diagonal lines).

As our model relies on translation-dependent OMM localization, many of the time scales are on the order of translation time (*τ*). Indeed, with this model, we are able to reproduce the rapid loss of transcripts at the OMM upon microtubule (k_leave_→k_leave_/2; k_return_→k_return_/4) or dynein perturbations (k_return_→k_return_/4) observed experimentally (Figure 3F), with stable cluster transcripts lost less quickly and remaining slightly more enriched at the OMM after ∼10-20 minutes (Figure 4C), compared to sensitive cluster transcripts which are lost quickly (t ∼ *τ*). The model suggests that this greater effect of dynein perturbation is because dynein provides retrograde transport to return RNAs to the mitochondria-rich perinuclear region, while microtubules also play a role in anterograde transport that removes RNAs to the periphery.

Furthermore, there is a coupling between the fraction of transcripts at the mitochondria (F_mito_) and the translational initiation rate. Notably, translation efficiency is proportional to the number of ribosomes bound to a transcript, which in turn is proportional to the translational initiation rate (Figure 4D). Increasing or decreasing translational efficiency can dramatically change the fraction of transcripts on the OMM, and vice versa. Indeed, the F_mito_ is quite sensitive to the translation rates involved, including initiation (Figure 4E) and elongation (<u>Figure S5A</u>), where a twofold decrease in initiation timescale (i.e. increased initiation rate) leads to doubling of F_mito_. Additionally, the rate of nascent peptide maturation influences F_mito_, with higher initiation rates and faster maturation of the encoded nascent peptide leading to increased F_mito_ (<u>Figure S5B</u>). Faster maturation is also associated with higher F_mito_ when the k_return_ (dynein) is high (<u>Figure S5C</u>) or when the k_return_ and k_leave_ are varied (<u>Figure S5D</u>). Notably, for our transcripts, we see no significant differences in translational speed (<u>Figure S5F</u>) between the stable, intermediate, or sensitive cluster transcripts. There is a slight difference in transcript lifetime (<u>Figure S5G</u>), and protein length (<u>Figure S5E</u>) between the stable and sensitive clusters but these differences do not contribute appreciably to the observed modeling outcomes.

## Discussion

Cells respond to external cues on timescales faster than transcription and RNA export, yet how cells can dynamically remodel their spatial RNA organization remains poorly understood. Here we leverage the spatial and temporal resolution of APEX-seq to quantify RNA localization dynamics at the OMM in living cells, and find that hundreds of OMM-proximal transcripts, including many nuclear-encoded mitochondrial mRNAs, require continuous active transport to maintain their localization. Acute disruption of microtubule-based transport, particularly dynein, rapidly remodels the OMM-localized transcriptome within minutes, without major changes in RNA abundance, mitochondrial positioning, or global translation. Time-resolved profiling reveals distinct kinetic classes of RNAs, and ribo-eCLIP measurements show that translation efficiency is a major predictor of transcript retention at the OMM. Integrating these data into a spatiotemporal kinetic model suggests that ribosome-mediated anchoring and motor-driven transport together determine RNA localization dynamics. These findings support a model in which RNA localization is not merely a static targeting mechanism, but an actively maintained regulatory state that enables cells to tune local translation on minute timescales.

The model reveals tight coupling between translation and mitochondrial RNA localization. Because the system responds on the translation timescale (*τ*), this coupling could provide rapid control of protein synthesis at membranes such as ERM^25^, OMM^46,47^ or lysosomes^140,141^ where local translation occurs. Notably, membranes such as ERM translate thousands of secretory proteins, including those presented to the cell surface in cancers, so these mechanisms apply to thousands of transcripts within cells. Intriguingly, the Mayr group has shown that the same transcript when forced to localize to the ERM versus the cytosol produces more protein^142^. In this study, we extend this conceptual framework to the OMM but add a kinetic dimension to the study. Our kinetics analysis reveals that there are mechanisms driven by ribosome anchoring and active transport that can dynamically control the fraction of RNAs on the mitochondrial surface, which in turn is correlated with a high translation initiation rate. These findings suggest the possibility that cells, perhaps in response to external stressors including viruses, cancer drugs or environmental cues, can respond within minutes by altering translation (increasing or decreasing) through localization. In many respects, such rapid alteration of translation is known to occur in condensates such as stress granules or P-bodies that are dynamic and able to sequester RNAs to suppress translation. However, translation at a membrane such as the OMM or ERM allows for, in principle, tunable and rapid increase or decrease of translation. Furthermore, while specific protein factors implementing such programs remain unknown, our modeling suggests that in principle, cells may be able to achieve either selective modulation of individual or small group of genes through binding of specific proteins that alter their membrane localization, or alternatively, systems-level changes in mRNAs translated on membranes through global changes^143,144^ in active transport, microtubule or actin stiffness, or recruitment or expression of translation factors (such as AKAP1^47,145^ and LARP4^48^ for mitochondria). The latter more global process would permit cells to modulate the translation of tens to hundreds of proteins, many of which might belong to large complexes with many proteins that need to change levels cooperatively. Depending on the exact cellular perturbation, both scenarios, specific or systems-wide control, might be desirable.

Finally, based on our modeling, it appears that there is a narrow range of translation initiation values correlating with maximal changes in localization, and this range contains the estimated typical initiation rate of mRNAs (Figure 4E). The model further indicates that this regulatory capacity is greatest within a restricted range of initiation and elongation rates, where the fraction on the mitochondria is tunable or responsive (Figure 4F), and not completely on (F_mito_ ∼1) or off (F_mito_ ∼ 0). This insight raises the possibility that in our system, and perhaps many others, RNA localization operates in a regime not necessarily optimized for efficiency, but for maximal translational responsiveness, which in turn might explain the widespread mRNA localization patterns in diverse systems and organisms. Thus, by performing out-of-equilibrium RNA localization measurements coupled with modeling, we raise the intriguing possibility that cells continuously spend motor energy to maintain localized mRNA pools at membranes, allowing translation to be tuned within minutes.

### Limitations

This work was performed in a single cell type (HEK) for which we have prior APEX-seq, and where recent proximity-specific ribosome profiling is revealing the regulation and logic of local mitochondrial translation^46,47^. In these cells mitochondria are perinuclear, and thus the strong dependence on the retrograde-motor dynein likely reflects this cellular organization, and other motors may dominate in cells with more polarized or peripheral mitochondria. A second limitation is that APEX-seq reports on proximity rather than binding, which confounds our interpretation as many likely reside at ER-mitochondria contacts. Third, the spatiotemporal model supports a tight coupling between RNA localization and translation, but the present study does not define the downstream consequences of mitochondrial protein production, protein import, or mitochondrial function. Finally, future studies will be needed to show how cells may use this kinetic coupling to respond more effectively to external perturbations.

## Author contributions

F.M.F. conceived the project and performed experiments. S.S. performed experiments, analyzed sequencing data, conducted imaging and image analysis, and generated the figures. S.N. performed machine learning-based feature assignment to clusters. M.E.R. wrote the pipeline for processing sequencing data. X.W. prepared samples for imaging. P.C. and A.I.B. developed the spatiotemporal model and performed biophysical simulations. J.Y.C. and E.L.V.N. performed ribo-eCLIP. T.T.T. and D.L.K. performed polysome profiling. F.M.F. and S.S. wrote the manuscript with input from other authors.

## Acknowledgements

F.M.F. acknowledges support from CPRIT (grant no. RR210012), NIH/NHGRI (grant no. R00HG010910), NIH/NIGMS (grant no. 1R35GM154922), Welch Foundation (grant no. Q-2186-20240404), Caroline Wiess Law seed fund for research in molecular medicine, and BCM seed funds. S.N. was supported by a Houston Area Molecular Biophysics (HAMBP) T32 GM150582 fellowship. F.M.F. and E.LV.N are CPRIT Scholars in Cancer Research. E.L.V.N. acknowledges support from NIH/NHGRI (R35HG011909). D.L.K. acknowledges support from NIH (grant no. 1R35GM137819). A.I.B. acknowledges support from a Natural Sciences and Engineering Research Council of Canada (NSERC) Discovery Grant, an NSERC Alliance International Catalyst Grant, start-up funds provided by the Toronto Metropolitan University Faculty of Science, by the Toronto Metropolitan University Faculty of Science Dean’s Research Fund, and computational resources provided by the Digital Research Alliance of Canada (alliancecan.ca). We thank Jitendra K. Meena, Devi P. Boggupalli, Noah Keogh, Brennan Keogh, and Kyle E. Nielson for helpful discussions to analyze imaging data. We thank members of the Fazal lab for providing feedback on the manuscript.

## Declaration of interests

D.L.K. is a named inventor on a patent owned by Houston Methodist Hospital related to the generation and use of circular RNA as a therapeutic platform. Further, D.L.K. has founded a company to commercialize that circular RNA technology and is a member of the scientific advisory board of different companies seeking to do the same. Both T.T.T. and D.L.K. are named inventors on invention disclosures filed with the Houston Methodist Hospital Office of Technology Transfer. E.L.V.N. is a co-founder, on the Scientific Advisory Board, equity holder, and paid consultant for Eclipse BioInnovations, on the SAB of RNAConnect, and is inventor of intellectual property owned by the University of California San Diego. E.L.V.N.’s interests have been reviewed and approved by the Baylor College of Medicine in accordance with its conflict-of-interest policies. All other authors declare no competing interests.

## Data availability

The datasets generated during this study have been deposited in the Gene Expression Omnibus (GEO). Drug perturbation and siRNA-treated datasets are available under accession number GSE329576. Nocodazole time-course and translation inhibitor datasets are available under accession number GSE116008.

The genome browser tracks can be found at:

https://genome.ucsc.edu/s/surbhi/motor perturbations tracks

https://genome.ucsc.edu/s/surbhi/Nocodazole time course

**Correspondence and requests for materials should be addressed** to Furqan M. Fazal.

## Methods

### Mammalian cell culture

HEK293T cells (<10 passages) expressing APEX2 targeted to the OMM and cytosol were cultured in a 1:1 DMEM/MEM mixture supplemented with 10% fetal bovine serum, 100 units/ml penicillin, and 100 µg/ml streptomycin at 37°C in a humidified chamber containing 5% CO_2_. U2OS cells were cultured under the same conditions. Cells were obtained from ATCC (catalog number CRL-3216 for HEK293T and HTB-96 for U2OS), and were checked to be free from mycoplasma contamination (Lonza MycoAlert PLUS detection kit).

### Treatment with inhibitors of motor transport and APEX labeling

The APEX labeling was performed as described previously^14,63^. Briefly, HEK293T cells expressing APEX2 at the OMM and in the cytosol were grown on fibronectin-coated 10 cm dishes. Prior to labeling with hydrogen peroxide, motor transport inhibitors were added. For MyoVin-1 (Sigma Aldrich, 475984), a final concentration of 30 μM was used for 10 min; for CK-666 (Sigma Aldrich, SML0006) a final concentration of 10 μM was used for 10 min; for nocodazole (Sigma Aldrich, M1404) a final concentration of 10 μM was used for 10 and 30 min. For ciliobrevin D (Sigma Aldrich, 250401), a final concentration of 50 μM was used for 30 min. For the 30 min timepoint, the motor-transport inhibitors were added along with media containing biotin-phenol.

### RNA extraction and enrichment of biotinylated RNA

The labeled (+H₂O₂) and unlabeled (−H₂O₂) control cells were scraped from 10 cm dishes using cell lifters, transferred to 1.5 ml RNase-free tubes, and spun at 300g for 4 min. The supernatant was removed, and RNA extraction was performed using the RNeasy Plus Mini Kit (QIAGEN) according to the manufacturer’s instructions, with a minor modification: the RW1 buffer was replaced with RWT for washing the RNA column. The extracted RNA was eluted in RNase-free water, and RNA integrity was confirmed using a Bioanalyzer (RIN ≥ 8.5). The enrichment of biotinylated RNAs was performed as described previously^14,63^ by using 25-30 μg of the collected RNA per replicate. The recovered biotin-RNA was purified using RNA clean and concentrator 5 kit (Zymo Research) and used for preparing sequencing libraries.

### Preparation of libraries for sequencing

RNA-seq libraries were generated from biotinylated RNA using Illumina TruSeq stranded mRNA preparation kit (polyA+ selection) with modifications as described here^63^. The prepared libraries were quality controlled and sequenced on the Illumina Hiseq 4000 or NovaSeqX Plus to an average depth of ∼30 million reads (paired end 2x75) per replicate.

### Sequencing data analysis

The sequencing data were mapped to the human genome (GRCh38.p13) and DESeq2-based differential expression analysis was performed to identify OMM-enriched transcripts relative to unlabeled controls as described here^63^. To identify transcripts localizing to the OMM, ratiometric normalization was performed by subtracting the log_2_FC changes obtained from cytosolic APEX2 from those of OMM-APEX2 for all reported log_2_FC, unless otherwise specified.

### siRNA-based knockdown of dynein motor complex

The knockdown of the dynein motor complex was performed using siRNAs (SMARTpool, Horizon Discovery) targeting ACTR1A (L-012074-00-0005), DYNC1H1 (L-006828-00-0005) and LIS1 (L-010330-00-0005). The non-targeting control pool (D-001810-10-20) was used as a control. Briefly, HEK293T cells expressing OMM-APEX2 were seeded at 30% confluency on fibronectin-coated 10 cm dishes. siRNAs were transfected at a final concentration of 20 nM using DharmaFECT transfection reagent in antibiotic-free media. 24 h post transfection, the media was replaced with antibiotic-containing media and cells were grown for a total of 48 h post transfection. APEX labeling was then performed as described above, followed by enrichment of biotinylated RNA using streptavidin beads, library preparation, and sequencing. Knockdown was validated at the RNA level by sequencing and at the protein level by Western blotting. Antibodies against ACTR1A and LIS1 (sc-374586, mouse, 1:200) were used, with actin (ab8227, rabbit, 1 µg/mL) as a loading control. For each sample, 25 µg of total protein was resolved on a NuPAGE 4-12% Bis-Tris gel, and proteins were detected using IR dye-conjugated secondary antibodies (680 and 800 nm). For siRNA knockdown experiments, log₂FC at the OMM was calculated. The log₂FC values for OMM-localized RNAs shown in Figure 2F correspond to 1,072 RNAs in control siRNA-treated samples, 1,033 RNAs in ACTR1A KD, 1,064 RNAs in LIS1 KD, and 374 RNAs in DYNC1H1 KD.

### Nocodazole time-course analysis

The nocodazole time-course data (3, 6, 9, and 30 min) derived from HEK293T cells expressing APEX2 at the OMM or in the cytosol was downloaded from GEO under accession number GSE116008. The sequencing data was analyzed as described above and cytosolic fold-changes were used for ratiometric normalization. Sequencing data generated under basal conditions was used as the 0 min time point. The −H₂O₂ controls from the 3 min time point were used to calculate enrichment for the 3- and 6-min time points, whereas the −H₂O₂ controls from the 30-min time point were used to calculate enrichment for the 9- and 30-min time points.

### Ribo-eCLIP

Ribosome-associated RNAs were identified by eCLIP of RPS2 (Chang JY, Van Nostrand EL, et al. *in preparation*). Briefly, cells were washed with PBS and UV crosslinked as previously described^146^. Cells were lysed in eCLIP lysis buffer, nucleic acids were fragmented with Turbo DNase and limited RNase I treatment, and immunoprecipitation was performed with anti-RPS2 antibody (Fortis Life Sciences). Dephosphorylation, RNA adapter ligation, SDS-PAGE electrophoresis, nitrocellulose membrane transfer, RNA extraction, reverse transcription, cDNA adapter ligation, and PCR amplification were performed as previously described^146^. Samples were sequenced on the Illumina NovaSeq XP platform.

### Immunofluorescence staining and fluorescence microscopy

For immunofluorescence, cells (HEK293T and U2OS) were grown on coverslips and fixed in 4% paraformaldehyde for 20 min at room temperature. Cells were washed with PBS three times followed by permeabilization with 0.5% Triton X-100 in PBS for 15 min followed by PBS wash three times. Blocking was performed using 10% goat serum (Thermo Fisher, 50062Z), cells were then incubated with primary antibodies (anti-TOMM20, D8T4N rabbit 1:200; anti-FLAG, F1804-1mg mouse 1:500; anti-Tubulin, sc-32293 mouse 1:250) diluted in antibody diluent (Thermo Fisher, 003218) at 4°C overnight. Cells were then washed with PBST (PBS with 0.05% Tween 20) twice, followed by incubation with secondary antibodies (1:1000) for 1 h at room temperature. After the incubation, cells were washed again with PBST twice and mounted with mounting media containing DAPI. Fluorescence confocal microscopy was performed with a Zeiss LSM900 microscope with 63x oil immersion objective.

### RNA FISH assay

RNAscope based Fluorescent *In situ* hybridization was used as per manufacturer’s recommendation to image OMM localization of candidate RNAs in U2OS cells. Briefly, 500,000 U2OS cells were seeded on coverslips in a six well plate prior to the experiment and allowed to grow for 24 h, followed by fixation in 4% formaldehyde and dehydration through graded ethanol. After rehydration, cells were permeabilized with Protease III. Hybridization was carried out using RNAscope probes targeting ACTB (310141-C2), MT-RNR1 (425961-C3), CPS1 (464981-C2), and IARS2 (1223781-C1), followed by signal amplification and fluorescent detection using the RNAscope™ Multiplex Fluorescence Detection Kit v2. The probe hybridization and the development of fluorescent signal were performed according to the instructions in RNAscope™ Multiplex Fluorescent Detection Kit v2 (323110). Fluorescence confocal microscopy was performed with a Zeiss LSM900 microscope with 63x oil immersion objective.

Images were acquired as z-stacks, and quantification of RNA association with the mitochondrial surface was performed using Imaris 11.0.0. Briefly, mitochondrial segmentation was carried out using the signal in the MT-RNR1 channel and RNAscope puncta were detected using the spot-detection module for each RNA candidate. OMM localization of the RNA was measured as the minimum distance between the centroid of each RNA spot and the mitochondrial surface.

### Sucrose gradient centrifugation for polysome analysis

Cells were treated with inhibitors of cellular transport as indicated above and fractionated using linear 10-50% sucrose gradients as described earlier^147^. Briefly, 10^7 cells/ml were washed twice with PBS containing Emetine dihydrochloride (100 µg*/*ml, EMD Millipore,324693-250MG) and 10 mM MgCl_2_. All further steps were carried out on ice using pre-chilled centrifuges. Cells were then scraped off the dish and pelleted by centrifugation at 2000 rpm for 3 min. Cell pellets were lysed using five volumes of ice cold lysis buffer containing 50 mM Tris-HCl, pH 7.5, 10 mM KCl, 10 mM MgCl_2_, 150 mM NaCl, 1% Triton X-100, 2 mM DTT, 0.5 mM PMSF, 100 µg*/*ml Emetine, Protease Inhibitor Cocktail (Millipore Sigma, P8340-5ML), Phosphatase Inhibitor II (Millipore Sigma, P5726-5ML), Phosphatase Inhibitor III (Millipore Sigma, P0044-5ML), and RNase-In Plus. The lysates were incubated on ice for 30 min with gentle mixing every 10 min, followed by centrifugation at 20000 rpm for 15 min to remove nuclei and other debris. The clear supernatants were then layered onto 10-50% linear sucrose gradients. These gradients were centrifuged for 3 h at 35000 rpm in a SW41 Ti rotor at 4°C. Fractions of 0.9 ml were collected from the bottom and the absorbance at 254 nm was continuously monitored using Brandel BR 188 Gradient fractionation system.

### Western blotting

Cell lysates were prepared by pelleting cells followed by washes with ice cold PBS and incubated on ice in M-PER lysis buffer supplemented with Protease Inhibitor Cocktail, Phosphatase Inhibitor II, Phosphatase Inhibitor III, and 0.05 mM PMSF. Proteins were separated by electrophoresis using pre-cast SDS-PAGE gel and transferred to a PVDF membrane. The membrane was blocked in 5% nonfat milk and incubated with primary antibody overnight at 4°C. The membranes were washed three times with 1x TBST and incubated with HRP-conjugated antibody for 2 h at room temperature. The protein bands were detected by western ECL Substrate, visualized using ChemiDoc MP Imaging System, and quantified by Biorad ImageLab software. We performed three independent replicates and representative blots are shown. Antibodies were used to detect eEF2 (Protein Tech, 20107-1-AP), eIF2α (Cell Signaling Technology, 5324S), phosphorylated eIF2α (Cell Signaling Technology, 3398S) and α-Tubulin (Protein Tech, 66031-1-Ig).

### Multiclass machine learning framework for cluster discrimination

To identify molecular features that best distinguish cluster identity, we implemented a supervised multiclass machine learning framework using scikit-learn (v1.3.0). Genes were assigned to one of three clusters: sensitive (N = 379), intermediate (N = 504), and stable (N = 189), and treated as independent observations. Cluster labels were used as the categorical target variable for model training.

Input features were automatically discovered from the input matrix and classified as numeric or categorical based on data type. Numeric features were median-imputed to account for missing values and standardized by Z-score normalization. Categorical features were imputed using the most frequent category and one-hot encoded, with unseen categories in the test set ignored during inference.

As one-hot encoding of categorical variables resulted in a high-dimensional feature space (894 total transformed features), we incorporated a feature selection step prior to model fitting. Feature selection was performed using SelectKBest with mutual information (mutual_info_classif) as the scoring function. Models were evaluated either without feature selection or with mutual information selection. The number of retained features (*k*) was treated as a hyperparameter and optimized jointly with model-specific parameters during cross-validation. Candidate values included 50, 100, 200, 400, 800, or all features.

We evaluated five supervised classification models: **(1)** Multinomial logistic regression with L2 regularization (SAGA solver)**. (2)** Multinomial logistic regression with L1 regularization (SAGA solver)**. (3)** Multinomial logistic regression with elastic net regularization (SAGA solver and tuned l1_ratio). **(4)** Random forest classifier **(5)** Histogram-based gradient boosted decision trees (HistGradientBoostingClassifier). To account for class imbalance, logistic regression models used inverse-frequency class weighting (class_weight = “balanced”), random forests used balanced subsampling (class_weight = “balanced_subsample”), and histogram-based gradient boosted trees used per-sample weights computed from balanced class weights and passed during fitting.

Data were split into training and test sets using an 80/20 stratified split with a fixed random seed. Hyperparameter optimization was performed exclusively on the training set using RandomizedSearchCV with stratified 5-fold cross-validation and macro-F1 score as the optimization metric. For each model, predefined hyperparameter distributions were explored using a fixed number of random draws (logistic regression: 40-60 iterations depending on penalty, random forest: 60, histogram gradient boosting: 40). The best-performing model for each architecture was refit on the full training set using the optimal hyperparameters. Final model performance was assessed on the held-out test set using accuracy, macro-F1, confusion matrices, precision/recall, and one-vs-rest ROC and precision-recall curves. Overfitting for each model was assessed by subtracting the test macro-F1 from the train macro-F1. To estimate performance stability beyond a single split, we performed repeated stratified cross-validation on the full dataset using RepeatedStratifiedKFold (5 folds x 10 repeats) and assessed each model’s macro-F1 mean and standard deviation across repeats.

For linear models, feature importance was quantified using fitted coefficients. Coefficients were reported per-class, and features were additionally ranked by mean absolute coefficient magnitude across classes. For tree-based models, feature importance was computed using Gini impurity (random forest only) and permutation importance. Permutation importance was evaluated on the test set using 30 random shuffles per feature.

### Stochastic simulations

The stochastic simulations couple an mRNA translation model to a cellular localization model within a Gillespie algorithm (Figure 4A&B). We apply an existing translation model^137^ which has each mRNA with L_mRNA_ amino acids, ribosomes that initiate translation at rate k_init_ if the first codon is not occupied by another ribosome, and ribosomes that step forward at rate k_elong_ if the next codon is not occupied by another ribosome (including ending translation by a ribosome at this rate when the ribosome is at the final codon). Ribosomes at or beyond L_mRNA_ of 100 amino acids can mature their nascent polypeptide at a rate k_maturation_ and mRNAs require at least one ribosome with a mature nascent polypeptide to associate with mitochondria and mRNAs are degraded at rate k_d_.

The translation parameters of this mRNA translation model are modified compared to previous work: We use translation initiation rates k_init_ in the range 0.025/s – 0.05/s, following a typical translation rate measured in mammalian cells (0.028/s – 0.056/s)^148^ and consistent with other translation initiation rate measurements in mammalian cells^149,150^, an elongation rate of 5 amino acids/s, approximately corresponding to measurements in mammalian cells^151,152^; a typical mRNA length of 600 amino acids, approximately corresponding to the median length of the genes we consider; and an mRNA degradation rate of (1/6000)/s, approximately corresponding to the mean cytoplasmic lifetime of genes^8^. An MTS maturation rate of 1/40/s was previously found to be consistent with measurements in yeast^137^; however many rates in mammals are slower than in yeast (e.g., the typical yeast translation initiation rate of 0.1/s^151^ is faster than the mammalian rate of about 0.03/s). Accordingly, the typical k_maturation_ we use is (1/100)/s.

The cellular localization model coarsely describes mRNA position in the cell as in the perinuclear region (N), the peripheral region (P), or associated with mitochondria (M). mRNAs that have newly emerged from the nucleus are located in N and can transition to other regions. mRNAs are transported from N to P at rate k_L_ and return to the N from P region at rate k_R_. As mitochondria are typically concentrated in the perinuclear region, mRNA in the perinuclear region N with at least one nascent polypeptide can associate with mitochondria at rate k_on_. mRNA associated with mitochondria that lose their last mature nascent polypeptide immediately end their mitochondrial association and return to the perinuclear state (this is represented in Figure 4A with k_off_). The dynamics of mRNA with a mature MTS can be described by

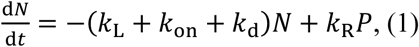

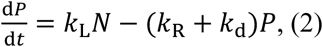

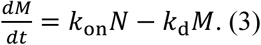

The dynamics of mRNA that do not have a mature nascent polypeptide can be described by

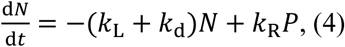

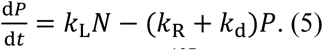

Previous work^137^ estimated the diffusive search time by an mRNA for mitochondria in yeast as approximately two to twelve seconds, depending on mitochondrial density. Since mammalian cells are larger than yeast cells, we use the longer end of this range, k_on_ = (1/10)/s. Motor-driven transport of cargos across distances of tens of micrometres that stochastically attach and detach from motors can require hundreds of seconds^153^, providing an estimate for transport times between the perinuclear region and the periphery. Diffusion will lead to faster departure from the perinuclear region than return from the peripheral region (there is more area or volume in the periphery of a circle or sphere than near the centre), and although motors can drive transport both away from and towards the perinuclear region, microtubules are denser near the nucleus than in the periphery. Accordingly, we use estimated values of the rate k_L_ = (3/200)/s to leave the perinuclear region for the periphery and rate k_R_ = (1/200)/s to return from the periphery to the perinuclear region.

The fraction associated with mitochondria is calculated as

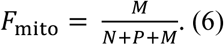

For steady-state calculation of F_mito_ (Figure 4E&F), many sample mRNA lifetimes are simulated. Each mRNA is associated with mitochondria for a fraction of its lifetime F_mito,i_. The average across many mRNA is calculated as

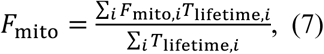

with T_lifetime,i_ the lifetime of each mRNA. For calculation of F_mito_ in transient scenarios that have not reached a steady state, which occur following a perturbation (Figure 4C&D), mRNA are considered to have left the nucleus at some time point up to several hours before the time of the perturbation until an hour following the perturbation (mRNA nuclear departures are uniformly selected in this time range). At the time of the perturbation and for the time points following the perturbation, the mRNAs that have left the nucleus but have not yet degraded (as the denominator) and the mRNAs that additionally are associated with the mitochondria (as the numerator) are used to determine the fraction of mRNA at that time that are associated with the mitochondria.

## Supplementary Figures

**Figure S1.**
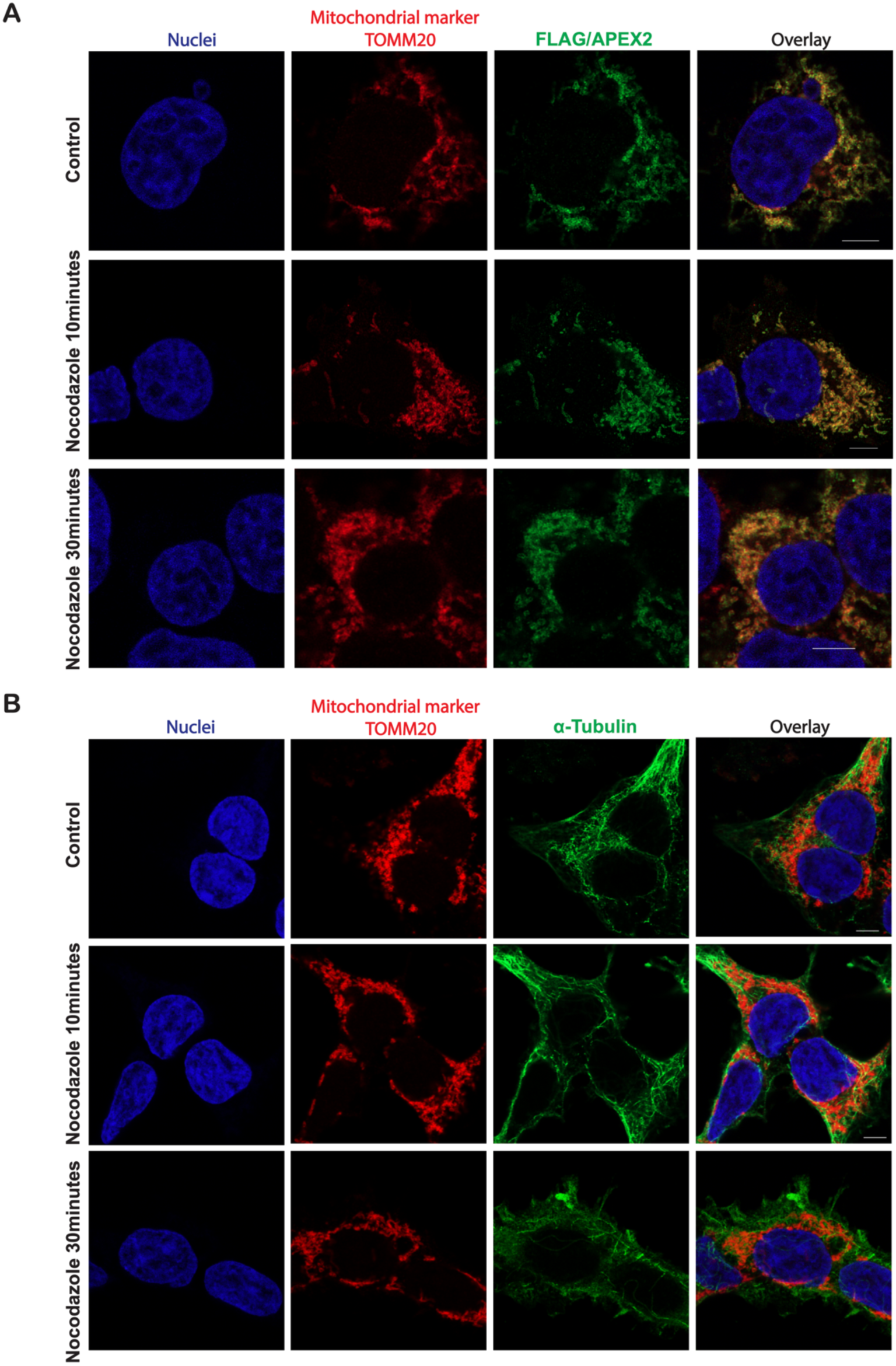
(A) Immunofluorescence images showing that APEX2 remains localized at the OMM in the presence of nocodazole, with co-localization of APEX2/FLAG and the mitochondrial marker TOMM20 in HEK293T cells, scale bars-5μm. (B) Immunofluorescence images depicting microtubule (α-tubulin) organization and mitochondrial distribution (TOMM20) in HEK293T cells treated with nocodazole for 10 and 30 minutes, scale bars-5μm.

**Figure S2.**
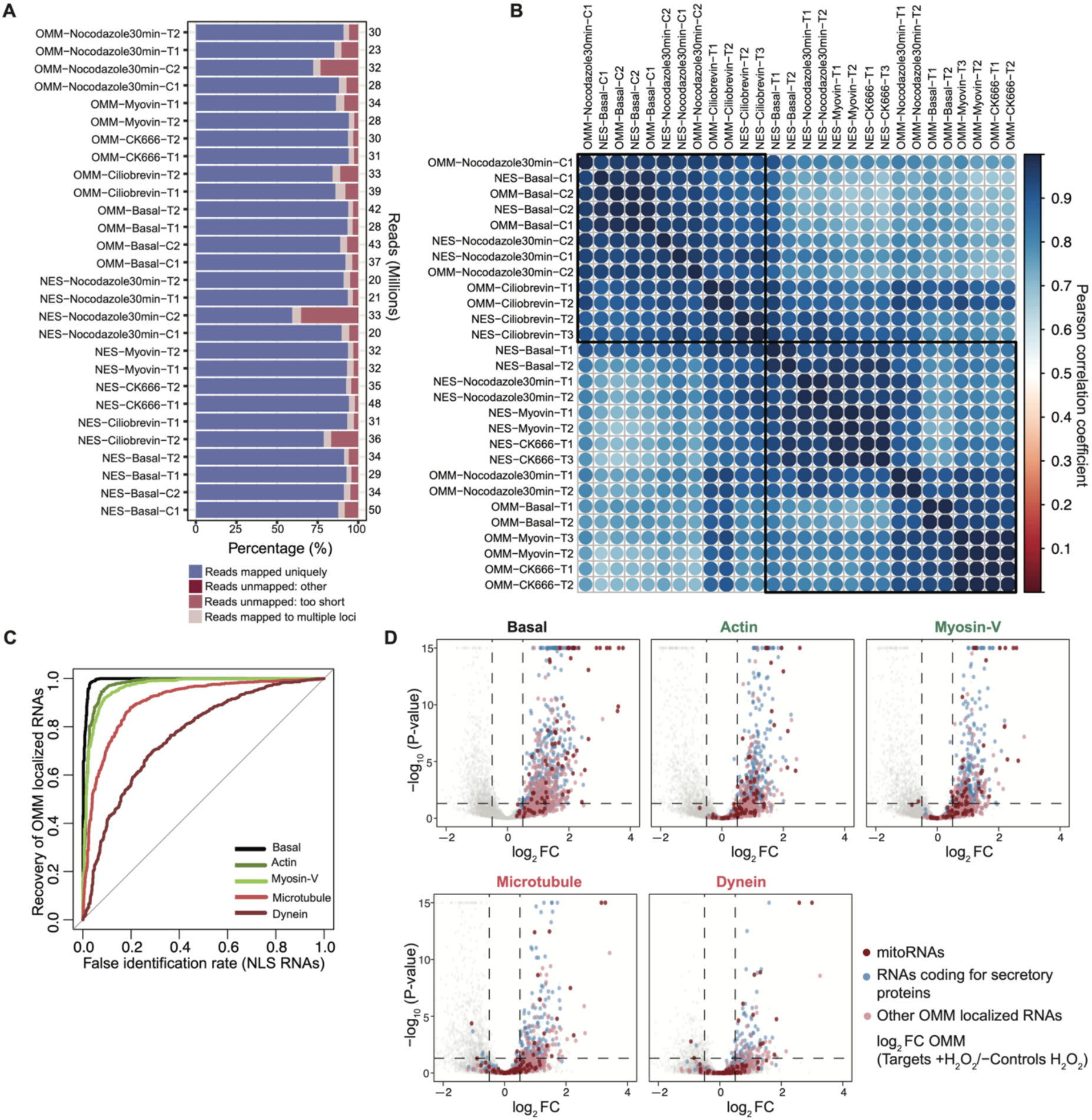
(A) Mapping statistics of APEX-seq libraries generated under basal and cellular transport perturbations. Bars represent the percentage of uniquely mapped reads and the total number of reads. (B) Pearson correlation coefficients of sequencing libraries, showing correlations among control (−H_2_O_2_) replicates across all conditions and among target (+H_2_O_2_) replicates within each condition. (C) ROC curves depicting the recovery of OMM localized mRNAs under basal and motor transport perturbations. The true-positive set consists of RNAs localized to the OMM under basal condition; the true-negative set contains RNAs that localize to the nucleus (NLS). (D) Volcano plots showing APEX-seq mediated enrichment of RNAs at the OMM (categorized as mitoRNAs, secretory, and others) under basal and cellular transport perturbation conditions.

**Figure S3.**
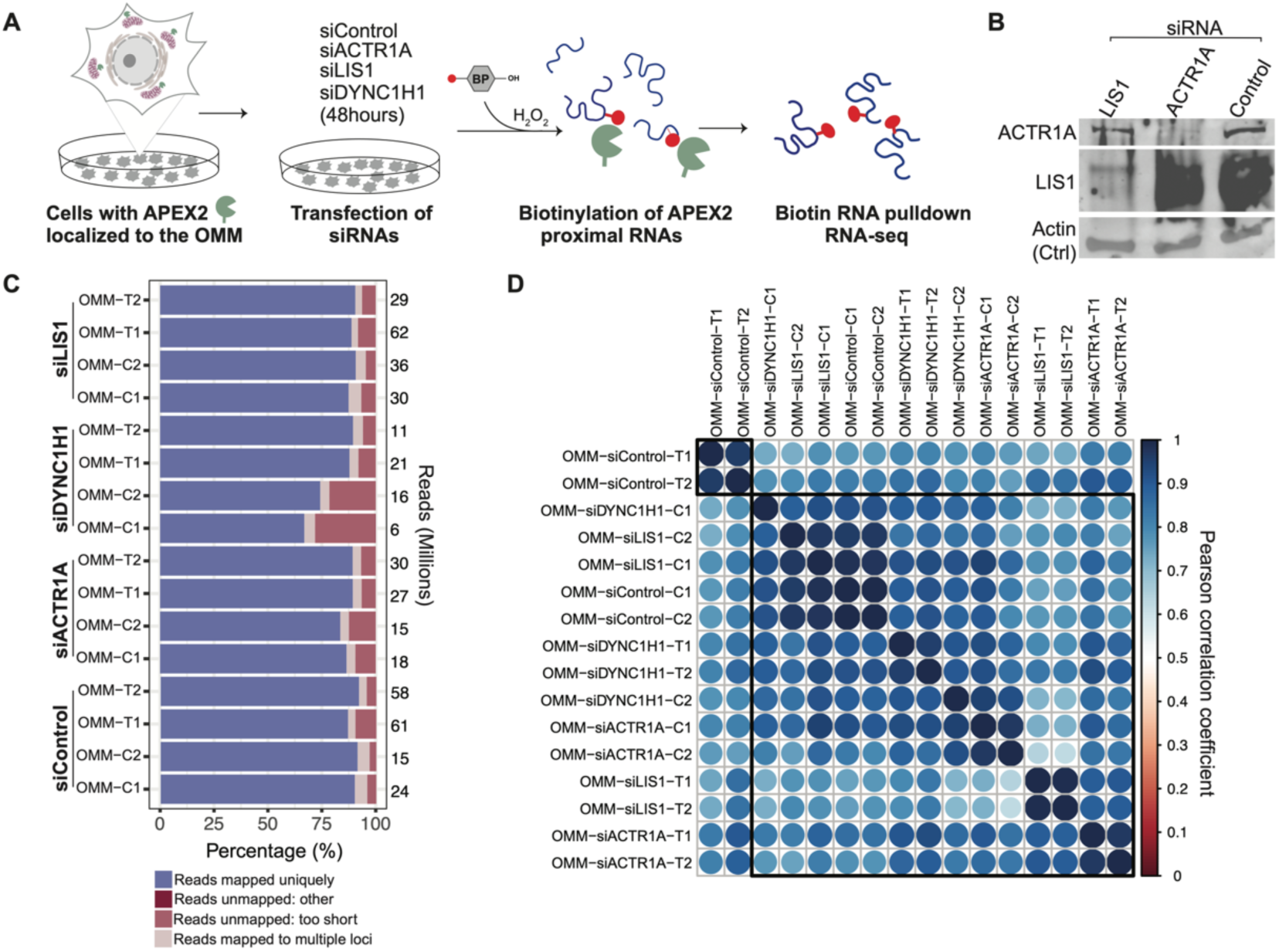
(A) Schematic of the experimental strategy for profiling RNAs at the OMM using APEX-seq after siRNA-mediated knockdown of the dynein transport complex. (B) Representative western blot showing knockdown of components of the dynein motor complex, β-actin used as loading control. (C) Mapping statistics of APEX-seq libraries from samples treated with siRNAs targeting components of the dynein motor complex. (D) Pearson correlation coefficients of sequencing libraries derived from siRNA-based knockdown of components of the dynein motor complex.

**Figure S4.**
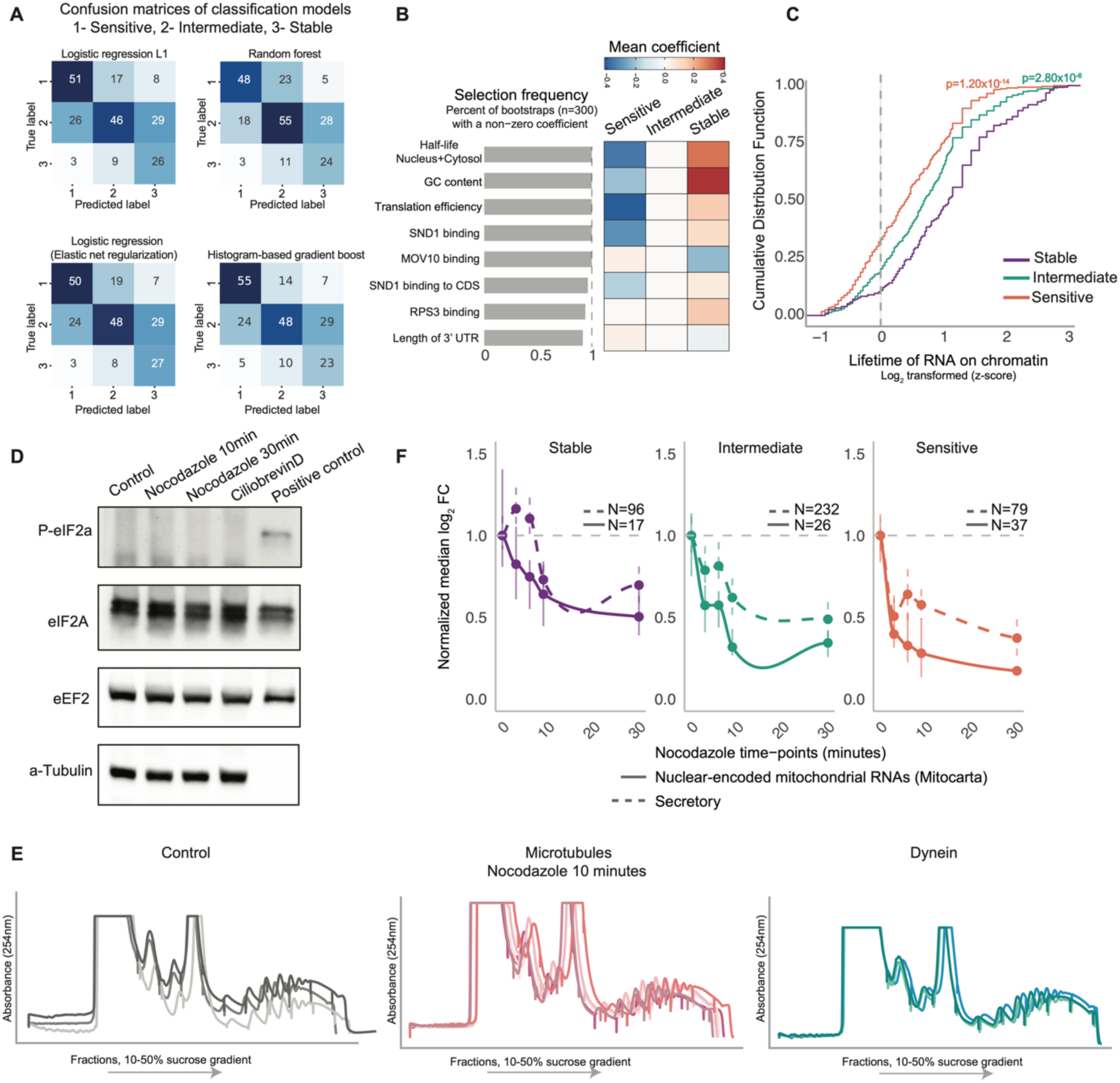
(A) Confusion matrices illustrating the performance of classification models used in this study. (B) Elastic-net regression analysis summarizing importance of top 8 features across 300 bootstraps. (C) Z-score-normalized RNA lifetimes on chromatin across clusters (stable, N = 160; intermediate, N = 401; sensitive, N = 299). (D) Western blot showing that the stress response is not activated by motor transport perturbations, as indicated by the absence of phosphorylated eIF2á. (E) Polysome profile traces of individual replicates on a 10-50% sucrose gradient from cells treated with inhibitors of the cellular transport. (F) Time-course profiling of OMM-localized transcripts (secretory and MitoCarta) across clusters following nocodazole treatment. Each point represents the cytosol-normalized median OMM localization value at each time point and the bars represent the interquartile range.

**Figure S5.**
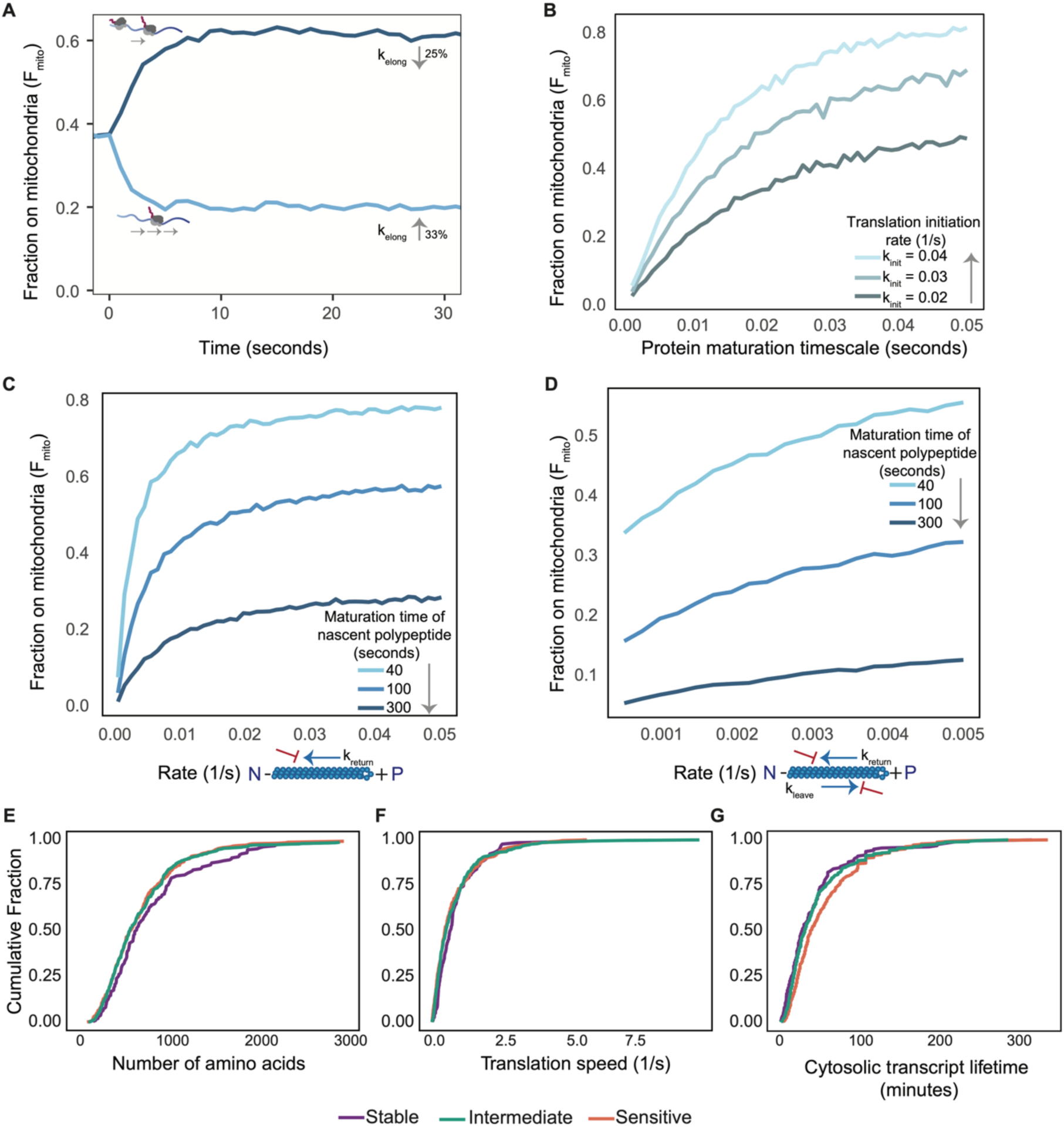
(A) Fraction F_mito_ of mRNA associated with mitochondria vs time since translation parameter perturbation. The curves show change in F_mito_ (for intermediate initiation rate) following increase of the elongation rate by 33% or decrease of the elongation rate by 25%. (B) F_mito_ vs maturation timescale for different initiation rates (k_init_ = 0.02-0.04/s). F_mito_ increases with maturation, with higher initiation rates yielding higher steady-state mitochondrial association. (C) Effect of varying the return rate to the perinuclear region K_return_ on F_mito_ for different MTS maturation times. (D) Effect of varying the rate of return to and departure fromt from the perinuclear region on F_mito_ for different MTS maturation times. (E) Cumulative fraction of the number of amino acids encoded by RNAs in each cluster, showing similar length distributions (stable, N = 182; intermediate, N = 487; sensitive, N = 374). (F) Cumulative fraction of translation speed for RNAs in each cluster, showing similar distributions of elongation step timescale (stable, N = 131; intermediate, N = 342; sensitive, N = 281). (G) Cumulative fraction of transcript cytosolic lifetime for RNAs in each cluster, showing similar cytosolic lifetimes across clusters (stable, N = 158; intermediate, N = 408; sensitive, N = 322).

